# Development of the Mouse and Human Cochlea at Single Cell Resolution

**DOI:** 10.1101/739680

**Authors:** Kevin Shengyang Yu, Stacey M. Frumm, Jason S. Park, Katharine Lee, Daniel M. Wong, Lauren Byrnes, Sarah M. Knox, Julie B. Sneddon, Aaron D. Tward

## Abstract

Hearing loss is a common and disabling condition, yet our understanding of the physiologic workings of the inner ear has been limited by longstanding difficulty characterizing the function and characteristics of the many diverse, fragile, and rare cell types in the cochlea. Using single-cell RNA-sequencing and a novel clustering algorithm, CellFindR, we created a developmental map of the mouse and human cochlea, identifying multiple previously undescribed cell types, progenitor populations, and predicted lineage relationships. We additionally associated the expression of known hearing loss genes to the cell types and developmental timepoints in which they are expressed. This work will serve as a resource for understanding the molecular basis of hearing and designing therapeutic approaches for hearing restoration.

## Introduction

Hearing loss is an inherent feature of human biology, with high frequency hearing loss typically starting at age 10 (Stelmachowicz et al., 1989). By age 75, half of all people will have disabling hearing loss (Stevens et al., 2013). The WHO estimates over 550 million people worldwide have disabling hearing loss. Despite the massive burden of hearing loss worldwide, our understanding of the normal biology of the human cochlea and how it goes awry in disease states is limited.

The common pathophysiology underlying the most frequent causes of hearing loss is dysfunction or loss of hair cells of the cochlea. Sound transmitted to the cochlea creates a pressure wave that depolarizes mechanotransductory hair cells and leads to the release of glutamate at the ribbon synapse, and thus the transmission of the signal via spiral ganglion neurons to the brainstem. These hair cells and neurons exist in stereotyped positions, within a highly ordered three-dimensional structure composed of many morphologically distinct cells.

Inner ear development begins with a thickening of the ectoderm, termed the otic placode (Kelley, 2006). The otic placode invaginates to ultimately form the epithelial lined otocyst. Within the otocyst Sox2-positive epithelial prosensory patches are specified, one of which will give rise to the cochlea. As the cochlear duct extends and curves, there is continued morphogenesis and differentiation of the cells lining the cochlear duct in a wave extending along the duct (Basch et al., 2016). The cells of the developing cochlear duct are organized into layers, with epithelial cells lining the lumen, periotic mesenchyme surrounding these cells, and ossifying bone surrounding the periotic mesenchyme (Fig 2d). Ultimately, the cochlea consists of three distinct fluid filled spaces, the scala vestibuli, scala media, and scala tympani. The hair cells and their supporting cells exist within the scala media, which also includes interdental cells, marginal cells, and Reissner’s membrane cells among others.

Some key functions of the hair cells and neurons are clear, but our understanding of the diverse cell types within the cochlea has been limited because the relative diversity of cells within each cochlea limits the utility of bulk characterization, and isolating specific cell types is made difficult by their small numbers and fragility. Although progress has been made in transcriptionally mapping the distinct cells of the cochlea (Burns et al., 2015; Durruthy-Durruthy et al., 2014; Ealy et al., 2016; Waldhaus et al., 2015), we do not have a global and systematic answer to the question of how many distinct cell types exist within the cochlea, at what point during development they are specified, or what distinct transcriptional programs they express (Basch et al., 2016). Furthermore, there have been no previously published single cell datasets on human cochleas, and is only limited bulk transcriptional information reported (Roccio et al., 2018; Schrauwen et al., 2016).

Developing such a map has important implications for human hearing loss. Although there are more than 150 genes implicated in human hearing loss (Shearer et al., 1993), in most cases we do not have a deep understanding of how they cause hearing loss, which cell types they affect and when they are actively being transcribed. Gene therapy approaches for genetic forms of hearing loss will depend on knowledge of the timing and localization of expression of the relevant targets. Regenerative medicine approaches to restore lost cells of the cochlea will depend on a deep understanding of the transcriptional events along the lineage trajectories of the relevant cell types. Finally, a detailed cellular map of the cochlea will serve as a crucial starting point for understanding disease states of the inner ear.

## Results

### Development of the CellfindR algorithm

We first aimed to identify distinct cell types in the developing mouse cochlea at E14.5 as this is a critical time point for specification of hair cells and their progenitors (Basch et al., 2016). We performed single-cell RNA-sequencing (scRNA-seq) on dissociated cochleas. In this experiment we identified 4,143 cells with an average of 1742 genes expressed per cell. Applying the Louvain clustering algorithm for community detection, we identified distinct epithelial populations representing the roof and floor of the cochlear duct, as well as multiple mesenchymal and neural crest populations (Fig 1b). Despite our ability to distinguish known subpopulations of cells based upon established markers, increasing the resolution parameter within the Louvain algorithm often failed to identify these subpopulations when analyzing the complete dataset. We noted that manually adjusting the resolution parameter of selected subsets of cells could identify biologically-relevant subpopulations. However, setting these parameters involves subjective judgement calls that could bias any analyses. We therefore developed an algorithm, CellFindR, that systematically and automatically performs automated and unbiased iterative subclustering to identify biologically significant groups (Fig 1a and **see Methods)**. Groups are labeled according to the iterative tree structure with main groups as the first integer and subsequent subgroups followed by a period, (i.e. 1.2.3).

**Figure 1:**
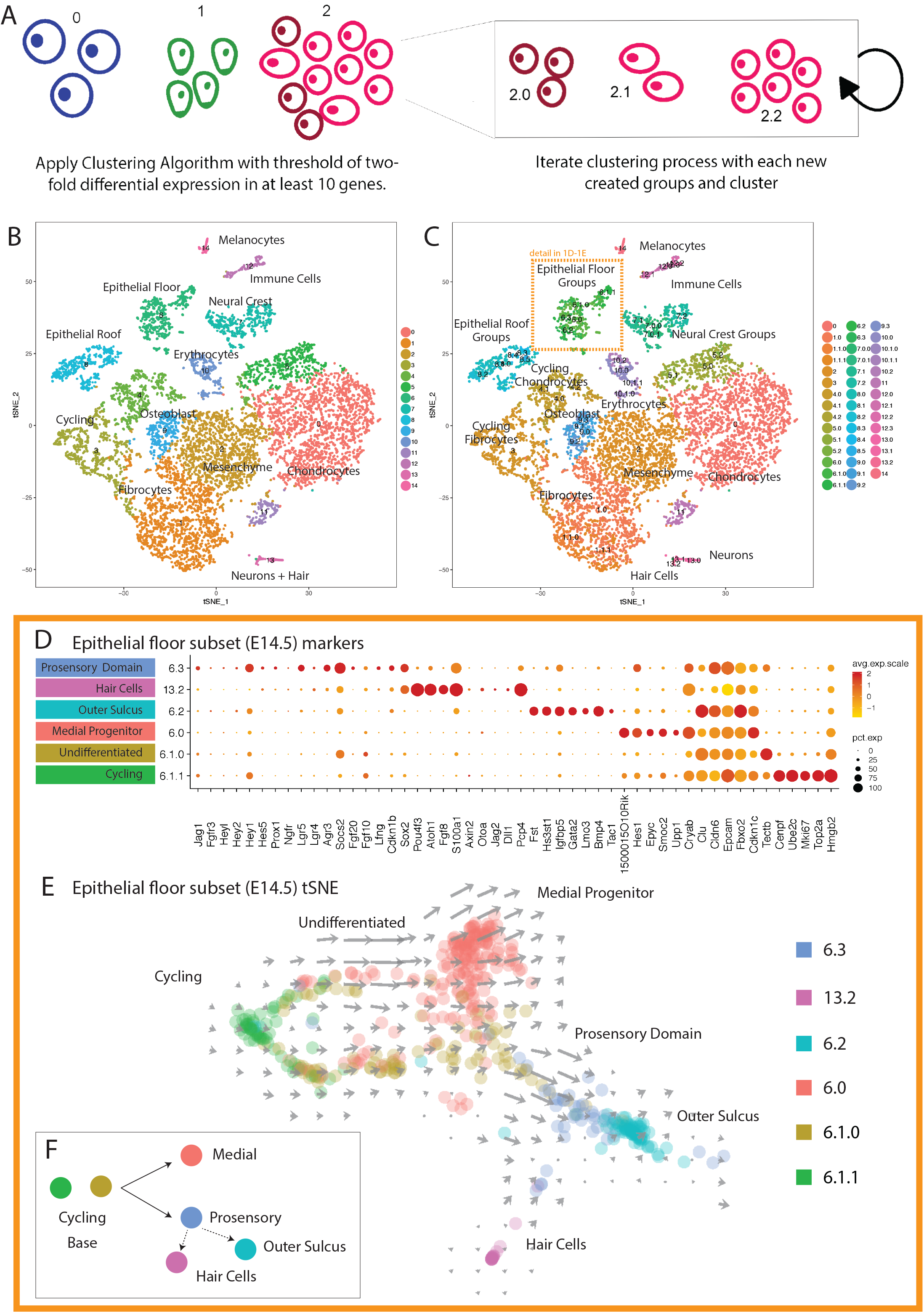
CellFindR methodology on single-cell RNA-sequencing data from E14.5 mouse cochlea. (A) Graphical representation of CellFindR algorithm, which uses iterative Louvain clustering with higher resolutions until the threshold of 10 genes with at least two-fold expression for each subcluster is violated. (B) t-SNE showing cell populations present in E14.5 mouse cochlea as identified by single Louvain clustering as per Seurat compared to (C) clusters generated by CellFindR. (D) Dotplot depicting subclusters of Group 6, the cochlear epithelium, showing diverse gene expression with known and novel markers of the prosensory, hair, outer sulcus, medial progenitor, undifferentiated and cycling cells. (E) Velocyto trajectory analysis on UMAP space showing progression of cycling and undifferentiated cells along with branching to medial, outer and prosensory domains. (F) Schematic of cochlear floor differentiaion at E14.5.

Applying CellFindR to the E14.5 dataset (Fig 1c), we discovered distinct cell types such as Schwann cells, melanocytes, multiple distinct subpopulations of fibrocytes, chondrocytes and distinct subgroups in the epithelium. CellFindR identified 44 distinct subpopulations in all, whereas our initial top-level Louvain clustering identified only 14 groups. In order to validate these computationally inferred subpopulations, we analyzed images of *in situ* hybridization performed on E14.5 mouse embryos from the Eurexpress database (Diez-Roux et al., 2011). Marker genes characteristic of the subpopulations stained distinct regions of the developing cochlea, supporting the validity of the CellFindR subgroups (Fig S1). Furthermore, CellFindR analysis of a biological replicate of E14.5 cochlea identified a similar number of subgroups with a similar pattern of gene expression (Fig S2). In order to validate that the CellfindR algorithm is generalizable to other datasets, we analyzed previously published human peripheral blood mononuclear cell and mouse brain scRNA-seq datasets from the 10X database. Within these datasets, CellFindR readily identified previously recognized subgroups of cells and additional distinct subgroups (Fig S3).

Based on Eurexpress data and marker genes, Group 6 define the epithelium of the cochlear floor, and Group 13.2 defines the hair cell population. In the cochlear floor epithelium, we identified five distinct cell states cycling, medial progenitor, (lateral) prosensory domain, outer sulcus, and an undifferentiated cluster of cells. (Fig 1d-f, **Table S1**). Velocyto (RNA velocity-based trajectory inference) analysis indicated a branched lineage trajectory whereby cycling progenitors exit the cell cycle and move toward the medial domain defined by expression of *Epyc, Hes1*, *Smoc2*, and *150015O10Rik* or toward the more lateral domain of the prosensory region cells defined by *Sox2, Eya1*, and *Fgfr3*, among other markers. The outer sulcus, anatomically lateral to the prosensory domain, is marked by expression of *Bmp4, Fst* and *Gata2*. This is consistent with previous reports of distinct cell fates of the medial and lateral domains (Ohyama et al., 2010).

### Developmental trajectories of the mouse inner ear

In order to generate a more comprehensive map of cochlear development, we performed scRNA-seq on E12.5, E14.5, E16.5, E18.5, and P2 mouse cochleas and analyzed each dataset individually using CellFindR. Proceeding from E12.5 to E18.5, CellFindR identified an increasing number of cell subgroups, which then plateaued at the E18.5 and P2 timepoints (Fig S4a). Concomitantly, the fraction of proliferating cells decreased across the arc of time (Fig S4b). Despite this decrease in proliferation, there continued to be large transcriptional shifts within the subpopulations after E18.5. These results are consistent with a model defined by a period of active specification and cell expansion terminating around E18.5, followed by continued differentiation at later timepoints.

Next, we batched the five datasets together and reanalyzed them as a single dataset with CellFindR in order to better understand which cell subgroups exist throughout development as well as to investigate their relationships between cell types across time (Fig 2a-b, d). We identified 172 distinct subgroups within this merged dataset. Of note, the total number of subgroups identified in the batched dataset was smaller than the cumulative number of subgroups identified without batching the data (306 groups), consistent with the biological expectation that some subgroups should exist across multiple developmental timepoints. Subgroups were always comprised of cells from contiguous timepoints (Fig 2c), thus validating CellFindR’s ability to identify cell subgroups that recapitulate biologically valid populations that exist in development. The development of the cochlear duct in a morphogenetic wave is reflected in our results as the presence of discrete developmental subgroups existing at multiple timepoints, with cells at distinct basal-apical positions being at different stages along the developmental gradient.

**Figure 2:**
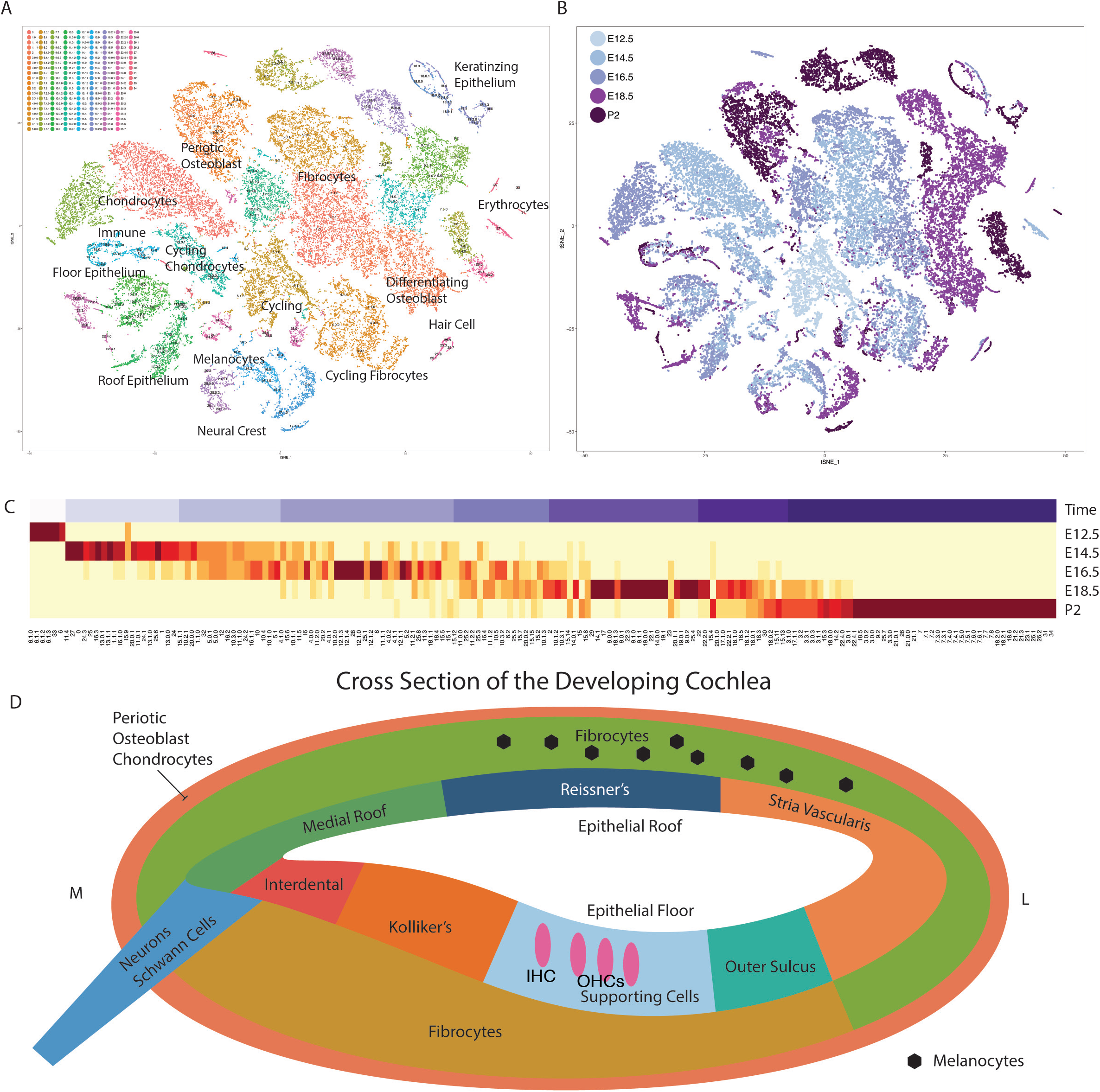
Lineage reconstruction of cell populations in the mouse embryonic inner ear across a developmental time course of E12.5 to P2 based on scRNAseq data. Batched embryonic mouse cochlea datasets in tSNE projection colored by (A) CellFindR groups and (B) the contribution by time point. (C) Heatmap showing contribution of each CellFindR group (in columns) by each time point (E12.5 to P2, in rows), ordered chronologically by weighted portion of the group, with color scale normalized across each column. A continuous purple scale representing the time course labels each CellFindR group with lightest representing mostly E12.5 and the darkest representing P2. (D) Schematic of locations of cells of developing cochlea at E18.5.

We further studied the epithelial subgroups of the cochlea, defined by *Epcam* expression. CellFindR identified four large classes of epithelial cell types, each of which was further subdivided into subclusters: cochlear floor, cochlear roof, hair cells, and keratin-expressing epithelium. We were able to infer the lineage relationships and developmental trajectories of each of these groups based on known marker genes, knowledge of which biological timepoints from which each CellFindR subgroup arose (purple gradient), and Velocyto-based trajectory inference analysis. This allowed us to generate a schematic illustrating the developmental transitions of the cochlear roof, cochlear floor, and hair cells (Fig 3).

**Figure 3:**
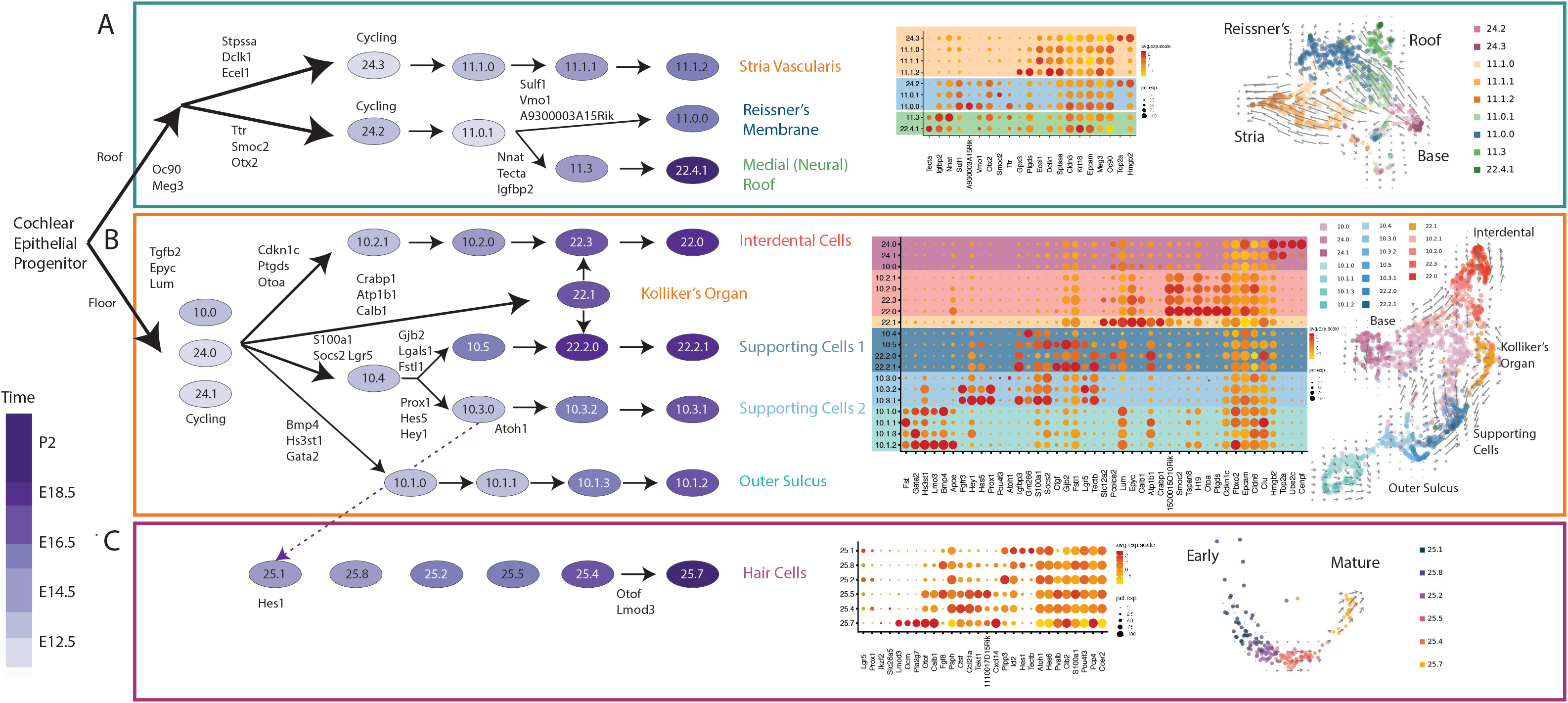
Lineage of Cochlear Epithelial and Hair cell Groups. (A-C) Schematic lineage trajectory of the cochlear epithelium showing progression from a predecessor to the (A) cochlear roof, (B) floor and (C) hair cells. Each sublineage is shown with representative dot plots and a supporting dimensional reduction figure (UMAP for roof and floor and PCA 1 vs 2 for hair cells) with velocyto overlay that aided in the construction of the lineage. Cochlear roof split into three groups the stria vascularis, Reisnner’s membrane and the neural medial portion. Cochlear floor split into groups representing interdental, Kolliker’s organ, supporting cells, and outer sulcus. Hair cells progress linearly along the time course with evolving markers as maturation occurs.

The cochlear roof cells (*Oc90*+), are transcriptionally distinct (Hartman et al., 2015) and decrease in relative number with maturation (Fig 3a),. This lineage first diverges into a branch that is destined to give rise to marginal cells of the stria vascularis (*Kcne1+*). Another divergence of this lineage gives rise to two distinct branches: *Vmo1+* cells of the Reissner’s membrane (Peters et al., 2007), and a *Tecta+ Nnat+* branch likely defining the medial portion of the cochlear roof (Freeman et al., 2015; Vendrell et al., 2015).

Among the cochlear floor cells (Fig 3b), a relatively undifferentiated *Tgfb2+ Epyc+* population (10.0) capable of proliferation (24.0, 24.1), can mature along a number of differentiation trajectories. One trajectory gives rise via several developmental intermediates to interdental cells (*Otoa+)* (Wan et al., 2013). Velocyto expression, shows *Smoc2* and *Pdgfc*, turn on in the undifferentiated population before subsequently turning off. The presence of the transient Kolliker’s organ in development is supported by our data (Dayaratne et al., 2014). Velocyto trajectory inference analysis shows this group (22.1) transitioning towards supporting and medial cells in the later timepoints. The *Bmp4+* lineage defines the outer sulcus, which is also characterized by the expression of *Fst, Gata2, and Lmo3* upon maturation (Ohyama et al., 2010). The progenitor cells of the cochlear floor (10.0, 24.0, and 24.1) express low, but detectable levels of *Lgr5*. *Lgr5* expression increases substantially concomitant with cell cycle exit and specification of the prosensory domain (10.4). The transition to the immediate bipotential hair cell progenitor (10.3.0) is best defined by upregulation of *Prox1* and the Notch effectors such as *Hes5* and *Hey1*. Within this progenitor, there is partial induction of *Atoh1* and *Pou4f3*, which either increase after specification of hair cell fate (25.1), or decrease if the cells follow a supporting cell trajectory (10.3.2, 10.3.1), concomitant with *Prox1* induction along this trajectory (Fekete et al., 1998; Kirjavainen et al., 2008).

Hair cells formed a separate group in our analysis, group 25, with contributions from all but one timepoint (no hair cells were isolated at E12.5) and seven developmentally distinct subclusters (Fig 3c). Velocyto analysis accurately ordered the subclusters in agreement with the known timepoints and prior biological knowledge of hair cell maturation from E14.5 to P2. At the latest time point, P2, maturation markers emerged, such as the outer hair markers *Slc26a5*, *Ocm* and *Calb1* (Scheffer et al., 2015). Orthogonally, hair cells were modeled with Monocle pseudotime analysis, which showed maturation along the time axis (Fig S5a-d). RNA velocity (Fig S5e,f) characterizes genes (*Otof)* actively turned on to be expressed at later timepoints, while *Ocm* is highly expressed at P2 only and stays on. *Fhl2, Flrt2, Pcdh15, Cdh23*, and *Myo15* are only on during earlier timepoints while *Tnc* and *Nme5* are turned on then off later. *Pvalb* is turned off early, but is subsequently turned on at P2.

The final epithelial group expresses multiple keratins and is comprised of middle ear mucosa with *Bpifa1* expression, mesothelium, and contaminant external ear canal skin (Fig S6). Neural crest derived cells such as melanocytes and Schwann cells, and their maturation, are also mapped in detail in the timecourse (Fig S7). The early Schwann cell marker *Npy* is expressed predominantly at early timepoints (E14.5-E16.5) group 20.0.1 and 16.3 (Ubink and Hökfelt, 2000) and later groups 20.1.0 and 17.1.1 show more mature markers: *Mbp, Mpz* (Liu et al., 2015). Given the presence of proliferation markers, these populations are actively dividing throughout this time period.

Mesenchymal populations make up a large proportion of the total cells in the merged dataset. The mesenchyme can broadly be separated into periotic (*Pou3f4+*) and outer otic capsule (*Pou3f4-)* populations (Phippard et al., 1998) (Fig 4). *Pou3f4* is known to regulate periotic mesenchymal populations and consistently demarcates these populations in our dataset. The diversity and anatomy of fibrocytes in the cochlea has been characterized previously, but known marker genes are often expressed in the multiple heterogeneous fibrocyte populations (Yoshida 2017). In our timecourse, we captured the progression of the periotic mesenchymal population to various fibrocyte populations (Fig S8), defined by marker genes such as *Vim* and *Col9a*, as well as unique subgroups with enriched expression of known deafness genes: *Coch*, *Otor*, and *Crym*. In addition, subgroups of *Pou3f4*+ cells expressing chondrogenic and osteogenic lineage markers including *Sox9*, *Mgp*, *Runx2*, *Dlx5*, and *Osx* were identified – these populations likely give rise to the osseous spiral lamina and modiolar bone (Rutkovskiy et al., 2016). In contrast, *Pou3f4*-mesenchymal cells are found in the outer otic capsule. In particular, we observe that *Pou3f4*-*Sox9*+ chondrocyte progenitors proliferate, differentiate, and mature during the timecourse of forming the outer otic capsule cartilage framework. These populations express common chondrocyte markers including *Col2a1*, *Matn1*, and *Hapln1* and decrease in relative number and proliferation over time. They are notably absent from the later timepoints (E18.5 and P2), when the framework is replaced by bone through the process of endochondral ossification by *Col1a1*+ osteogenic lineage cells.

**Figure 4:**
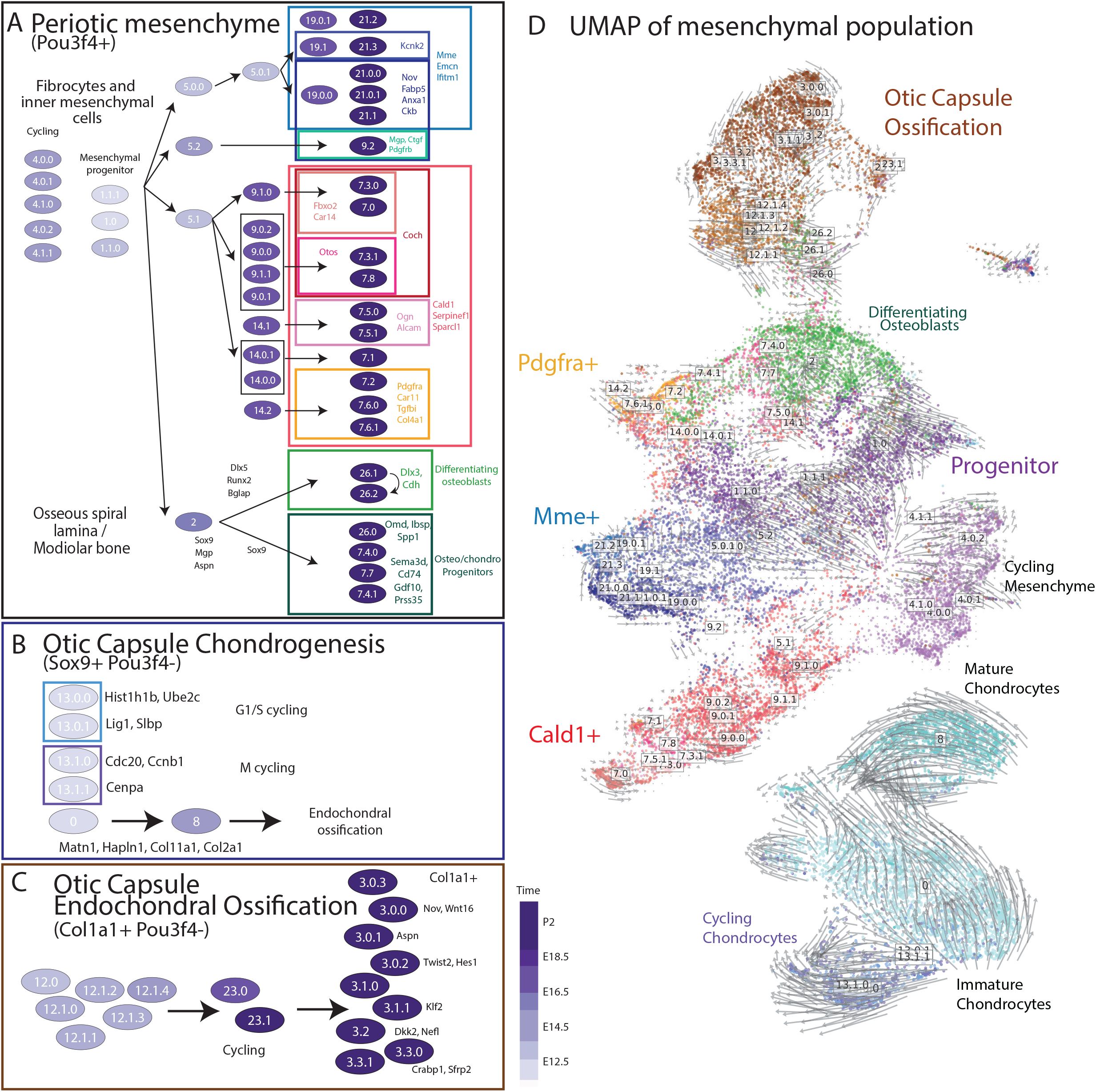
Mesenchymal lineages in the developing cochlea. Split into periotic and outer otic populations, (A) the periotic mesenchyme, marked by *Pou3f4* positivity, splits into pop fibrocyte groups and differentiating osteoblasts. The otic capsule (B) chondrocyte popoluation is cycling and ultimately is lost through endochondral ossification while (C) osteoblasts from the ossification process are also captured. (D) UMAP of the entire mesenchymal population with RNA velocity overlay.

Populations of immune and erythrocytes are also captured in the dataset (Fig S9). Multiple erythrocyte groups are captured, separated by age. Various myeloid and lymphoid cells are detected with velocyto analysis showing little to no velocities, consistent with the notion that these cells differentiated elsewhere then circulate or migrate to the cochlea.

### Adult mouse inner ear

In order to characterize the terminal and mature state of cells within the cochlea, we performed scRNA-seq on adult mouse cochleas. Mouse cochleas were dissociated and cells were sorted for Epcam positive and negative fractions. Using CellFindR, we identified 31 distinct subgroups within the Epcam-positive population and 49 groups in the Epcam-negative population. When added together, the number of groups is similar to that observed at P2. Hair cells and supporting cells of the Organ of Corti are distinguished along with cells of the inner and outer sulcus, marginal cells of the stria vascularis, and melanocytes along with mesenchymal and fibrocyte population (Fig 5a-b). In order to validate these inferred subgroups, we performed scRNA-seq on biological replicates of Epcam-sorted adult cochlea cells. When CellfindR was run independently on these samples, the inferred subgroups were remarkably similar to their cognate biological replicate as defined by fractional identity of top 100 most highly expressed genes in each group (Fig 5e-f). We used RNAscope staining to further validate our subgroup inferences. For example, fibrocyte subgroup 8.2, marked by *Otos* expression, localized predominantly to the spiral limbus. *Kcnk2* is specific for the 8.0 subgroup, which is defined by staining of the inner ridge medial to the spiral limbus. *Ppp1r1b* is expressed in fibrocytes of the stria vascularis, the spiral limbus and in the Boettcher cells and found in Epcam-negative groups 8.0, 8.1, 8.2. *Gpx3* marks marginal epithelial cells of the stria vascularis Epcam-positive groups 1 and 2 and also Epcam-positive group 6 representing Reissner’s membrane. Typically, we found novel subgroups of cells were often not marked by unique markers (such as *Kcnk2* above), but rather by unique combinations of markers which alone do not uniquely define subpopulations.

**Figure 5:**
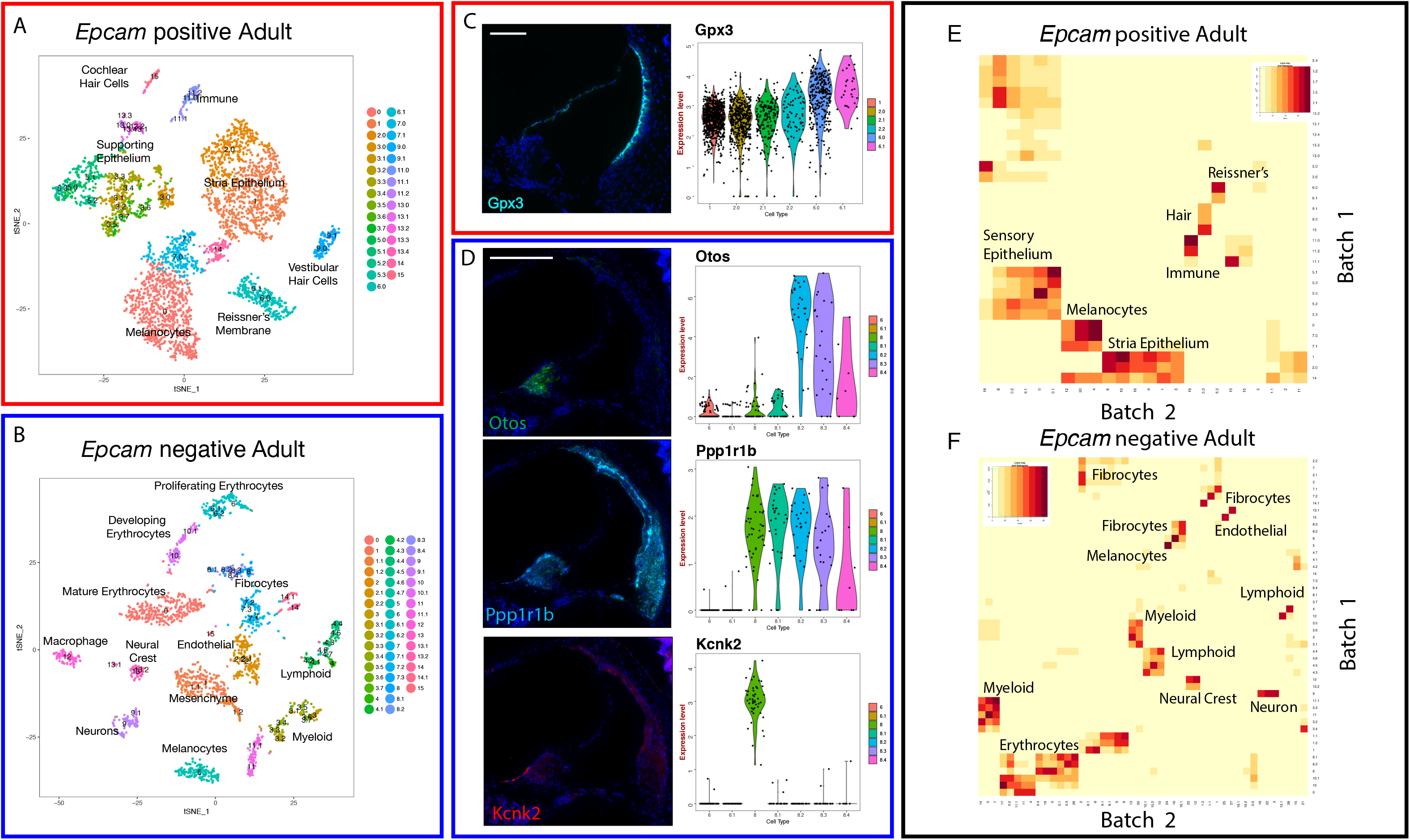
Adult mouse inner ear data. (A-B) tSNE representation of adult (A) *Epcam* enriched and (B) *Epcam* negative inner ear populations with labeled clusters of CellFindR groups. (C) RNAscope for *Gpx3*, marking marginal cells in the stria vascularis, and Reissner’s membrane (Groups 6.0 and 6.1). (D) RNAscope for *Otos* marks fibrocytes of the the spiral limbus (group 8.2). *Ppp1r1b* marks a majority of the fibrocyte population of the spiral ligament, stria vascularis and Boettcher cells which encompasses all of group 8. *Kcnk2* stains specific fibrocytes connecting the spiral limbus to Reissner’s membrane represented by group 8.0. Scale bars = 100 um. Biological replicates (Batch 1 and Batch 2) of adult (E) *Epcam* sorted positive and (F) *Epcam* negative are compared through their CellFindR groups, measuring the intersection of their top 100 expressed genes. Regions of high concordance are labeled with their cell identities.

Group 15 defines the adult cochlear hair cells (63 cells). At this resolution, we disproportionately captured inner hair cells over outer hair cells. CellFindR was unable to distinguish these groups due to the low numbers of outer hair cells. Supervised subgrouping using known markers *Slc26a5* (Zheng et al., 2000) and *Ocm* (Simmons et al., 2010), clearly distinguished the inner vs. outer hair cells. We were able to identify other differentially expressed markers such as *Strip2*, *Chrna10*, *Aqp11* for outer hair cells as well as the known inner hair cell marker *Slc17a8*/VGLUT3 (Seal et al., 2008) and additional novel markers inner hair cell markers such as *Igfbp5*, *Atp2a3*, and *Dnajc5b* (Fig S10).

### Human inner ear development

To identify how similar cochlear differentiation is between mice and humans, scRNA-seq was performed on human fetal inner ears at gestational age of 15, 17 and 23 weeks. These timepoints were selected as they span critical time periods of human cochlear specification. The 15 week and 23 week dissociated cochleas were *EPCAM-*sorted prior to scRNA-seq, while the 17 week cells were not sorted prior to scRNA-seq. CellFindR clustering was conducted for each experiment independently. For these human datasets, the overall architecture of the cells was remarkably close to the cell populations identified in the mouse cochlea. We identified similar populations of mesenchyme, cartilage, epithelium, neural crest and developing hair cells (Fig S11).

The epithelium of the floor of the cochlea in human shows a similar developmental trajectory to the mouse, with distinct populations corresponding to the key branches corresponding to the inner sulcus, interdental cells, supporting cells of the Organ of Corti, and the outer sulcus. In the 15 week dataset, Group 8 defines the interdental cells and markers, while markers for the four distinct regions are all present in a subset of the epithelium including cells that are *BMP4* positive (outer sulcus), *EPYC* positive (inner sulcus), and a few subgroups expressing supporting group homologs of genes from the mouse data. For 17 week data, group 6.1 expresses interdental markers and groups 6.4 and 6.5 expresses *BMP4* and other outer sulcus markers. At 23 weeks, group 3.1 shows outer sulcus markers while other groups showed less concordance with the mouse dataset.

In the human, as in the mouse, hair cells appear to arise from *LGR5*+ progenitors and show a decrease in *NOTCH* signaling and increase in *ATOH1* and *POU4F3* following specification. Hair cells are distinct groups for the 15 week data (group 11) and 17 week data (group 10) while for the 23 week data, four hairs cells are found in Group 16. In similar fashion to the mouse data, at 15 weeks we observe a low level of *ATOH1* expression in the presumptive bipotential progenitor cells (week 15 group 10), but by week 17 and definitely by 23 weeks we no longer observe a similar partially specified hair cell progenitor (Fig 6a-c). Comparing the maximally differentially expressed genes across Monocle pseudotime from the mouse hair cell trajectory to these developing human hair cells, there is remarkable similarity in the pattern of expression within maturing hair cells between mice and humans (Fig 6d). Following the levels of these marker genes, we can extrapolate that 15 week human hair cells correspond roughly to E16.5 mouse hair cells, 17 week human hair cells correspond to E18.5 mouse hair cells, and 23 week human hair cells correspond to past the P2 time point.

**Figure 6:**
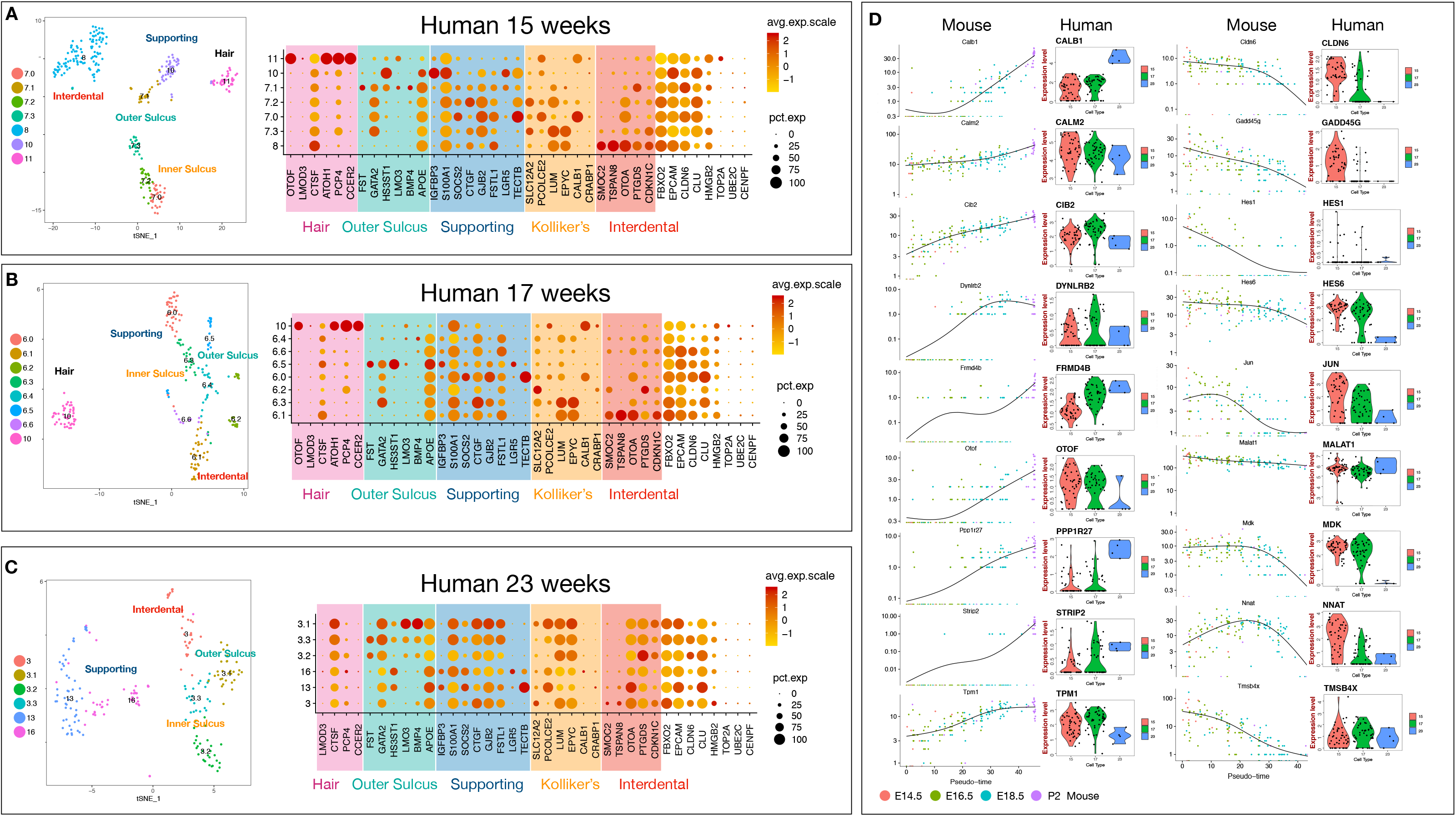
Gene expression patterns of fetal human (15, 17, 23 gestational weeks) hair cells and other epithelial cells compared to mouse. (A) (left) t-SNE of the epithelial subclusters of 15 week human fetal cochlea with associated dotplot (right)subdivided by CellFindR subgroups using genes of human homologs of mouse markers derived from the mouse time course data. (B) and (C) represent the same analysis as 15 weeks for 17 and 23 weeks, respectively. (D) Panel of top differential genes over the pseudotime trajectory generated by monocle analysis are plotted for mouse hair cells over E14.5 to P2 with comparative violin plots of human homologs for fetal hair cells from 15, 17 and 23 weeks. Upregulated genes over time are in the first column and downregulated genes are in the second column.

Other cell types exhibit similar trends in comparing human to mouse data. Oc90/OC90, similarly defines epithelial roof, disappears in the human population mostly by 17 weeks while in mouse, this group is decreasing at P2. Examination of epithelial marker genes and their expressions over time in mouse and in human reveal similar trends however, we observe that *BMP4* a strong marker for mouse outer sulcus is not highly expressed in the human sample (Fig S12). Otherwise, within the technical limitations of the human data, although the timing of events differed, we did not observe major differences between the transcriptional lineage trajectories of the mouse and human cochlea.

### Deafness gene mapping

Our transcriptional map was used to gain insight into the timing and expression pattern of genes implicated in hearing loss. A list of known hearing loss-causing genes was generated and the relative expression of these genes in the developing cochlea over time was plotted, normalized to expression in the maximally expressed CellFindR subgroup for each gene at each time point (Fig 7a). Some genes are maximally expressed early in development, including those that result in morphological abnormalities of the inner ear, such as *Chd7* (CHARGE syndrome) and *Six1* (Branchio-Oto-Renal). Other genes were expressed throughout development, such as *Pou4f3* (DFNA15) and *Cib2* (DNFB48, Usher type 1). Additional genes were expressed exclusively in adult cochlea, such as *Tmc1* (DFNA36) and *Whrn* (Usher type 2).

**Figure 7:**
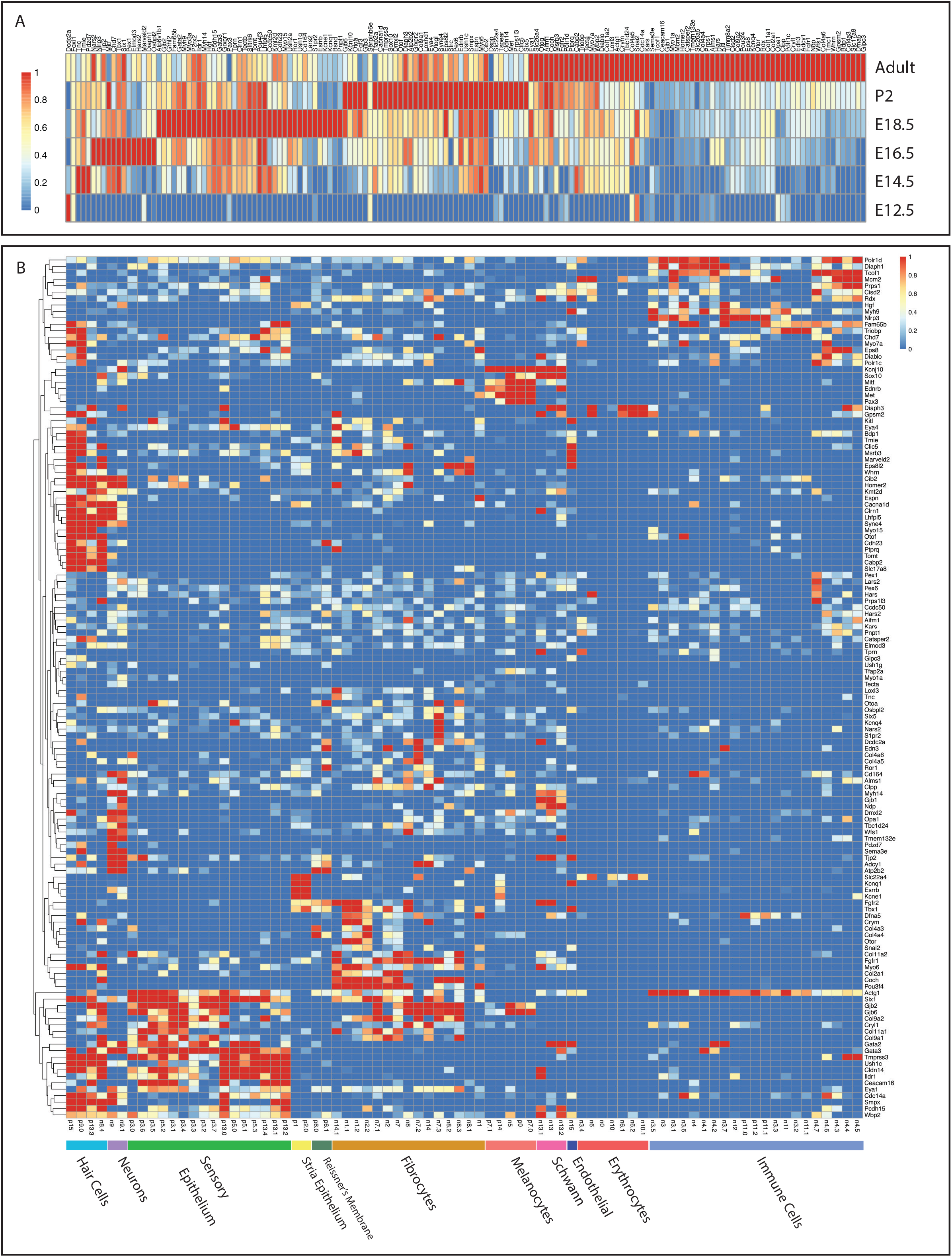
Deafness genes mapped across mouse development. (A) Expression levels of each deafness gene (columns) across all samples from E12.5 to P2 (rows), where expression is taken from the highest CellFindR group from that time point. The genes are sorted chronologically by highest expression from left to right and data are log normalized to 1 for each gene. (B) Expression heatmap of deafness genes across both the adult mouse *Epcam* sorted positive and negative datasets using CellFindR groups. Expression of all groups were combined together before being binned to a maximum of one and then hierarchically clustered together. Groups from the Epcam-positive dataset are prefixed with a “p,” while groups from the Epcam-negative dataset are prefixed with an “n”. Cell types are labeled for the column groups at the bottom.

In order to gain insight into the mechanisms by which these genes may cause hearing loss, we mapped the set of hearing loss genes onto our adult cochlea dataset. Many, but not all, hearing loss genes were expressed predominantly in the hair cells (Fig 7b). However, there were many clusters of genes that were not expressed in hair cells, but were instead expressed in other distinct groups of cells. For example, *Coch* is expressed within a subset of fibrocytes that also express *Pou3f4, Col9a2, and Col9a1*. On the basis of this expression pattern, we may predict, for example, that *Pou3f4* may be an important regulator of the effector function of this population that includes secretion of matrix components including *Coch* as well as collagens. Marginal cells express *Kcne1, Slc22a4, Esrrb*, and *Bsnd.* Furthermore, the same analysis was conducted to examine cell types for both the mouse embryonic and human fetal time course which identified both distinct cell type and developmental regulation of hearing loss gene expression (Fig S13, S14). In conclusion, based upon the diversity of timing and cellular tropism, there exists a diversity of mechanisms of genetic sensorineural hearing loss.

## Discussion

### CellFindR for subgroup identification

The identification of biologically relevant subgroups of cells within large single cell datasets is fundamental to the utility of recently developed single cell sequencing technologies (Kiselev et al., 2019). One key limitation of a number of implementations of clustering algorithms for scRNA-seq analysis is the number of parameters that need to be manually set, such as the resolution parameter in Louvain community detection-based clustering. It has therefore become common to perform subclustering analyses on groups of interest to identify more subtle differences between subpopulations within a group. However, it is difficult to know *a priori* which cells to include or exclude in particular subclusters. Combined, these issues introduce non-systematic biases that limit the utility of these algorithms, particularly for previously undescribed cell states. CellFindR presents a relatively simple automated solution to these issues and permits the identification of biologically valid subgroups of cells with minimal manual setting of parameters. We believe the reason why CellFindR is better able to identify subgroups of cells is related to the dimensionality reduction necessary at each step of subclustering. With a very heterogeneous group of cells, differential gene expression is going to be primarily driven by differences between major cell types, such as epithelium versus mesenchyme. Thus, more subtle differences between cell types that drive the identities of more closely related cell states are drowned out.

### A global view of cochlear development

The cochlear map developed in this study was constructed in a largely unbiased fashion. In the areas where there is deep prior biological knowledge, for example in hair cell development, the derived map is in excellent agreement with previous findings. Thus, we feel confident that the inferred subpopulations and lineage relationships reported here where there is less detailed knowledge (such as in Reissner’s membrane or mesenchymal populations) are likely to be biologically valid.

Based on the map developed here, cochlear development appears to be globally divided into three phases. First, there is a proliferative phase of relatively undifferentiated cells that rapidly declines after E14.5. Then, from E14.5 to E18.5 there is a specification phase where multiple distinct lineages are specified and the cochlea is largely populated by progenitor cells with little proliferation. This phase is then followed by a maturation phase from E18.5 into postnatal life where specified cells continue to alter their gene expression profiles, but retain the identities assigned during the specification phase. These transitions largely mirror previously published data regarding the patterning and differentiation of the prosensory domain, but this dataset extends these findings to additional lineages within both the epithelium and non-epithelial components of the cochlea.

Of particular interest are the distinct fibrocyte populations, all of which appear to be derived from *Pou3f4*+ periotic mesenchyme progenitors. Although their gene expression profiles are less distinct from each other than some of the epithelial populations, there appears to be a number of distinct cell subpopulations. With the exception of *Pou3f4* itself, the majority of hearing loss genes expressed within these groups encode for secreted matrix proteins such as collagens and cochlin which remain expressed in adulthood, indicating a likely ongoing role of expression of these genes in normal hearing. Distinct populations of cells also express high levels of aquaporins and distinct carbonic anhydrase genes such as *Car2, Car3, and Car14* which may indicate a role in ion and pH maintenance.

### Human cochlear development

Because of limited availability of primary samples and limitations of dissociation, our human fetal datasets were more sparse than our mouse datasets. Nonetheless, our analysis revealed remarkable similarity between human and mouse cell cochlear subpopulations. The timescale of the transitions in the differentiation of the human cochlea is extended relative to the mouse. Whereas mouse otic proliferation and specification is complete within one week, this process is apparently spread across many weeks in the human. We otherwise observed a remarkably similar pattern of key markers of distinct subpopulations in the developing human cochlea to the developing mouse cochlea. Thus, we believe that despite the size and timing differences of cochlear development between the mouse and human, the mouse is likely a very good model of human cochlear development. Indeed, the similarity in hearing phenotypes between mice and humans with similar hearing loss mutations is consistent with this notion (Chatterjee and Lufkin, 2011).

### Implications for regeneration approaches

There is currently great interest in restoring hearing to humans with sensorineural hearing loss using a variety of regenerative approaches. Although we observed *Lgr5+* supporting cells within the adult cochlea, we did not observe any resident cell populations in the adult cochlea that closely resemble the bipotential hair cell progenitors during their native differentiation trajectory. The lack of a similar immediate progenitor to hair cells may help explain the lack of regeneration in the mammalian cochlea under physiological circumstances. Further, we do not observe much proliferation within the immediate hair cell progenitors; rather, proliferation seems to be restricted to an earlier cochlear floor progenitor cell. This may help explain why current genetic and small molecule approaches to inducing regeneration within the adult cochlea require modulation of multiple signaling pathways prior to seeing new hair cell formation (Kuo et al., 2015; McLean et al., 2017). Because the immediate hair cell progenitors no longer exist, first a signal to place cells into a more progenitor like state is required, followed by Notch inhibition to guide differentiation down a hair cell fate. However, if such reprogramming is possible, it may be the case that the proliferative progenitors may be able to give rise to multiple supporting cell types in addition to hair cells. This may be advantageous in situations where the damaged sensory epithelium lacks not only hair cells but also the surrounding supporting cells (Devare et al., 2018).

### Implications for gene therapy

A number of proof of concept studies demonstrating the feasibility of virally mediated gene therapy in mouse models of monogenic hearing loss have recently been published (Akil et al., 2019; Al-Moyed et al., 2019; Nist-Lund et al., 2019; Pan et al., 2017). It is likely that our earliest gene therapy approaches are best targeted to those genes expressed predominantly in the mature cochlea, rather than earlier in development or throughout development. It is unsurprising, for example, that gene therapy in mice for *Otof, Tmc1, and Slc17a8* were all successful given that all of these genes are expressed almost exclusively in the mature cochlea (Fig 5b). Conversely, caution should be exercised in considering gene therapy for earlier expressed genes such as *Chd7* or *Six1* in children or adults with hearing loss. Furthermore, the distinct cellular tropism of hearing loss gene expression needs to be accounted for when selecting viral vectors and appropriate promoters for gene therapy. Although some currently available viral vectors appear relatively efficient at transducing hair cells (György et al., 2019; Landegger et al., 2017; Yoshimura et al., 2018), we will likely need to identify additional viral vectors capable of efficiently transducing other distinct cell types within the cochlea for those genes expressed within other cell populations.

## Acknowledgements

The authors would like to acknowledge Angie Koo and Heydi Malave for technical assistance. The authors would also like to thank Eunice Wan and Diya Vaka from the UCSF Human Genetics core, as well as the UCSF Flow Cytometry Core. The authors would like to thank Karl Koehler for helpful discussions. This work was supported by grants from the American Otological Society (A.D.T), Hearing Research Inc. (A.D.T.), the Program in Breakthrough Biological Research (A.D.T. and J.B.S.)., and the UCSF Resource Allocation Program (J.B.S.). K.S.Y. was supported by a grant from the UCSF School of Medicine, S.M.F. was supported by a fellowship from the American Otological Society and the UCSF Medical Scientist Training Program (T32GM007618), D.M.W. was supported by a National Science Foundation Graduate Research Fellowship and the Diabetes, Endocrinology, and Metabolism Training Program (T32 DK007418). L.E.B. was supported by an Achievement Rewards for College Scientists Foundation Scholar award.

## Author Contributions

Conceptualization, K.S.Y. and A.D.T.; Methodology, K.S.Y., A.D.T., J.S.P., S.M.F., D.M.W.; Software, K.S.Y. and J.S.P.; Validation, K.S.Y., A.D.T; Formal Analysis, K.S.Y., A.D.T., S.M.F., J.S.P., K.L.; Investigation, K.S.Y., S.M.F., K.L., D.M.W., L.B.; Resources, A.D.T. and S.M.K., S.M.F.; Data Curation, K.S.Y., S.M.F., J.S.P., K.L., A.D.T.; Writing – Original Draft, K.S.Y, A.D.T, J.S.P, S.M.F, K.L; Writing – Review & Editing, K.S.Y., S.M.F, J.S.P., S.M.K, J.B.S., A.D.T, Visualization, K.S.Y., S.M.F., J.S.P., K.L.; Supervision, S.M.K., J.B.S., A.D.T.; Project Administration, A.D.T.; Funding Acquisition, A.D.T.

---

**Figure S1:**
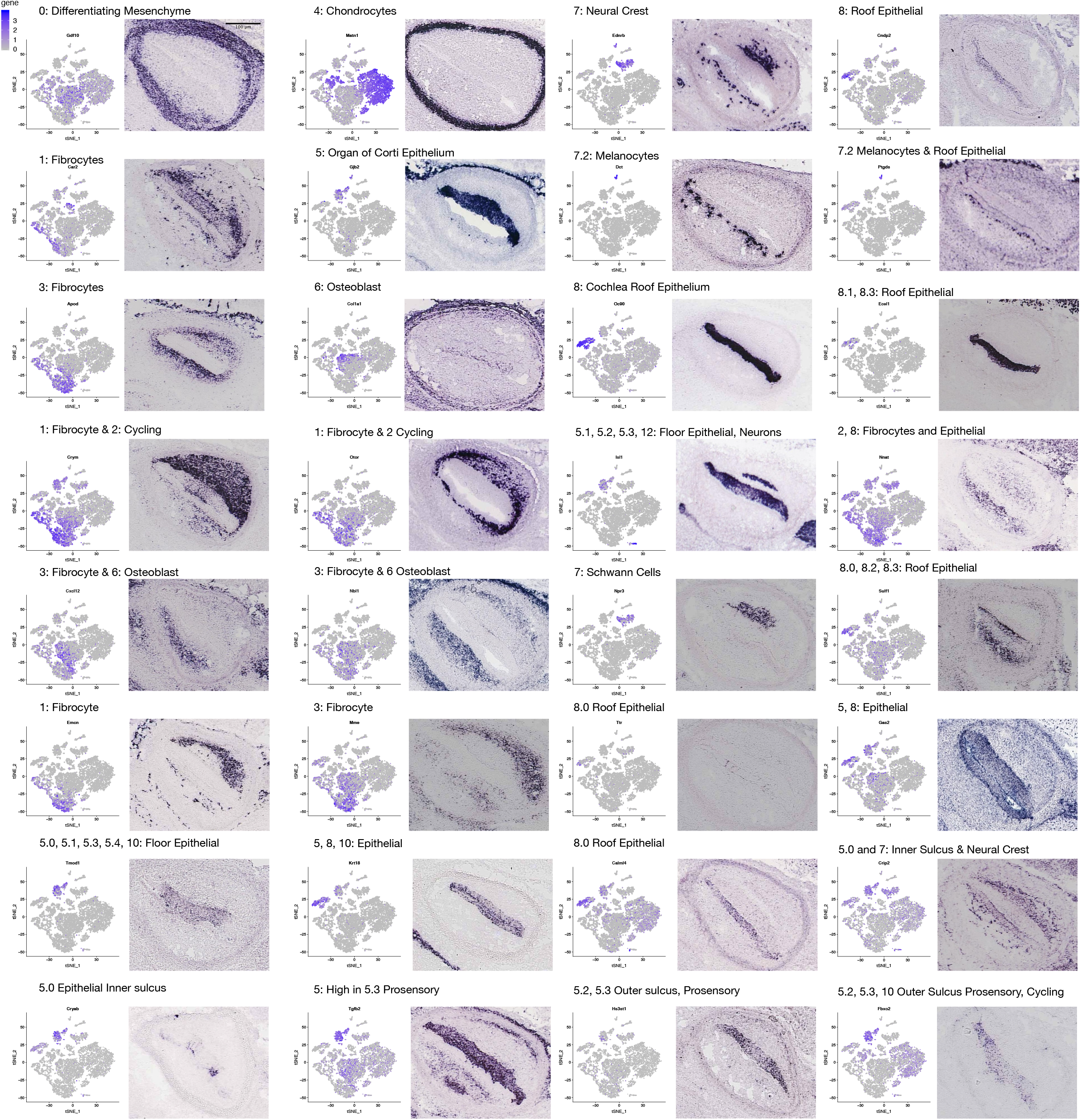
Validation of E14.5 markers through Eurexpress in-situ hybridization database. Identifying markers for E14.5 CellFindR groups confirmed via Eurexpress dataset. Full slice images were taken from the database and a 250 pixel by 200 pixel rectangle of a cross section was taken. Each gene is also accompanied by the tSNE expression plot from the E14.5 mouse dataset, showing specificity of single cell analysis and their correlation with canonical staining results. Scale bar = 100um.

**Figure S2:**
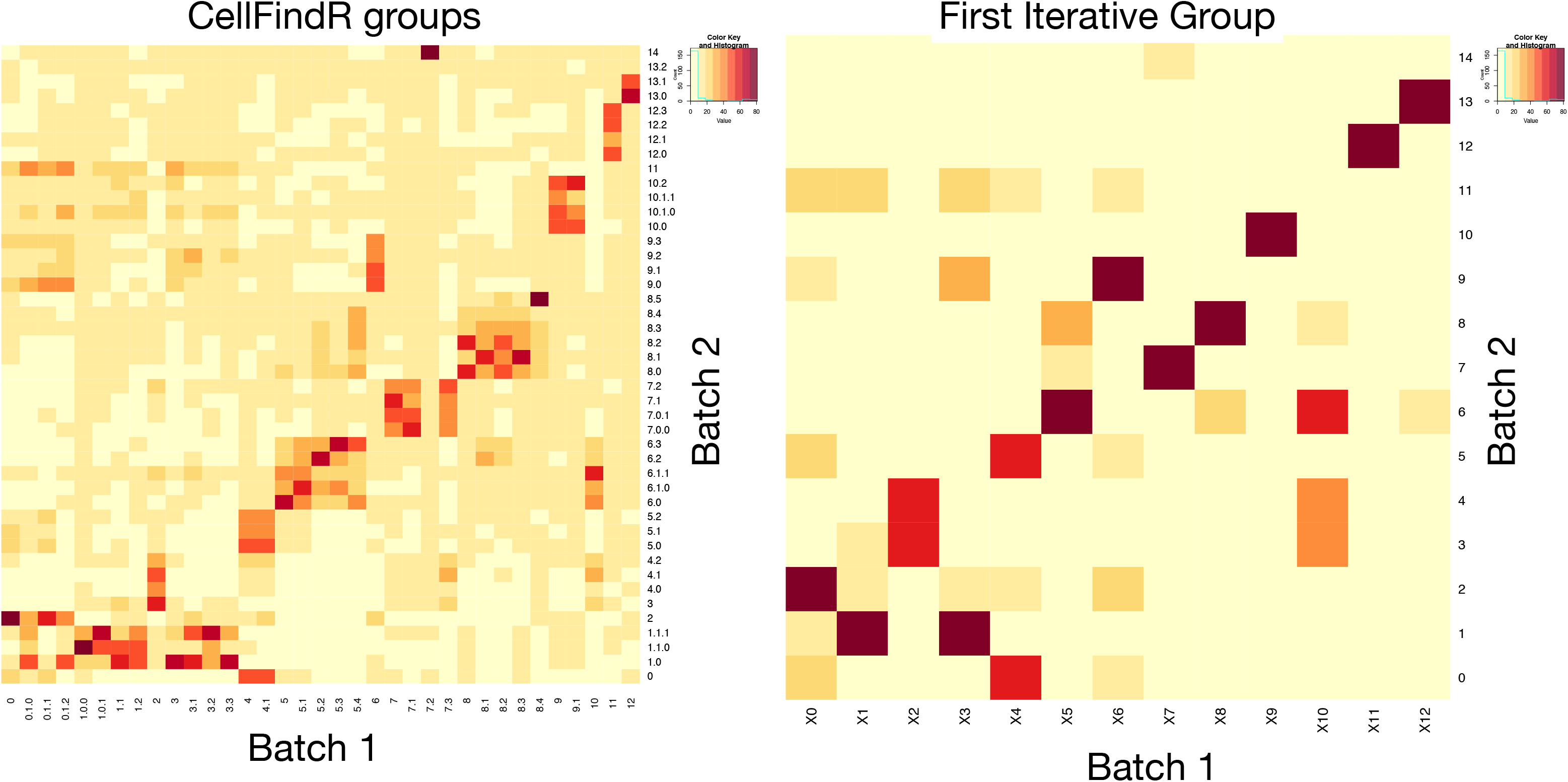
Correlation matrix of E14.5 mouse biological duplicates. Concordance of the intersection of the top 100 genes by expression of the (A) CellFindR groups and (B) first iterative layer of the louvain clustering groups compared between the E14.5 biological replicates.

**Figure S3:**
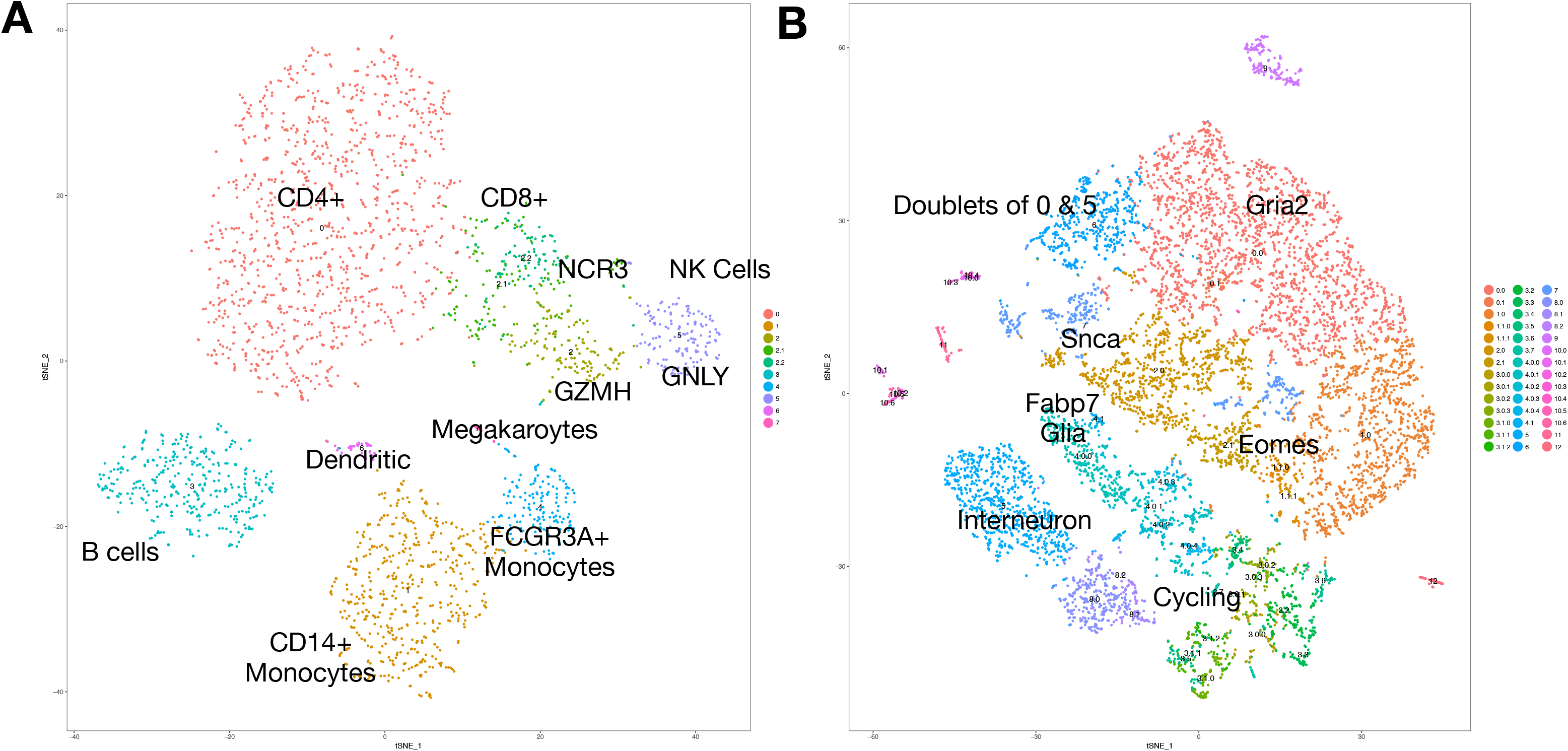
CellFindR algorithm on PMBC 2.7k from healthy human donor and brain 9k mouse E18 data. Both datasets (A) human pbmc, and (B) mouse brain, from the 10x Genomics website, were run with CellFindR algorithm and extracted more detail with subgroups beyond louvain clustering. For the pbmc data, specificity was extracted along the NK cells while in the mouse brain, primary cycling groups were able to be separated into phases.

**Figure S4:**
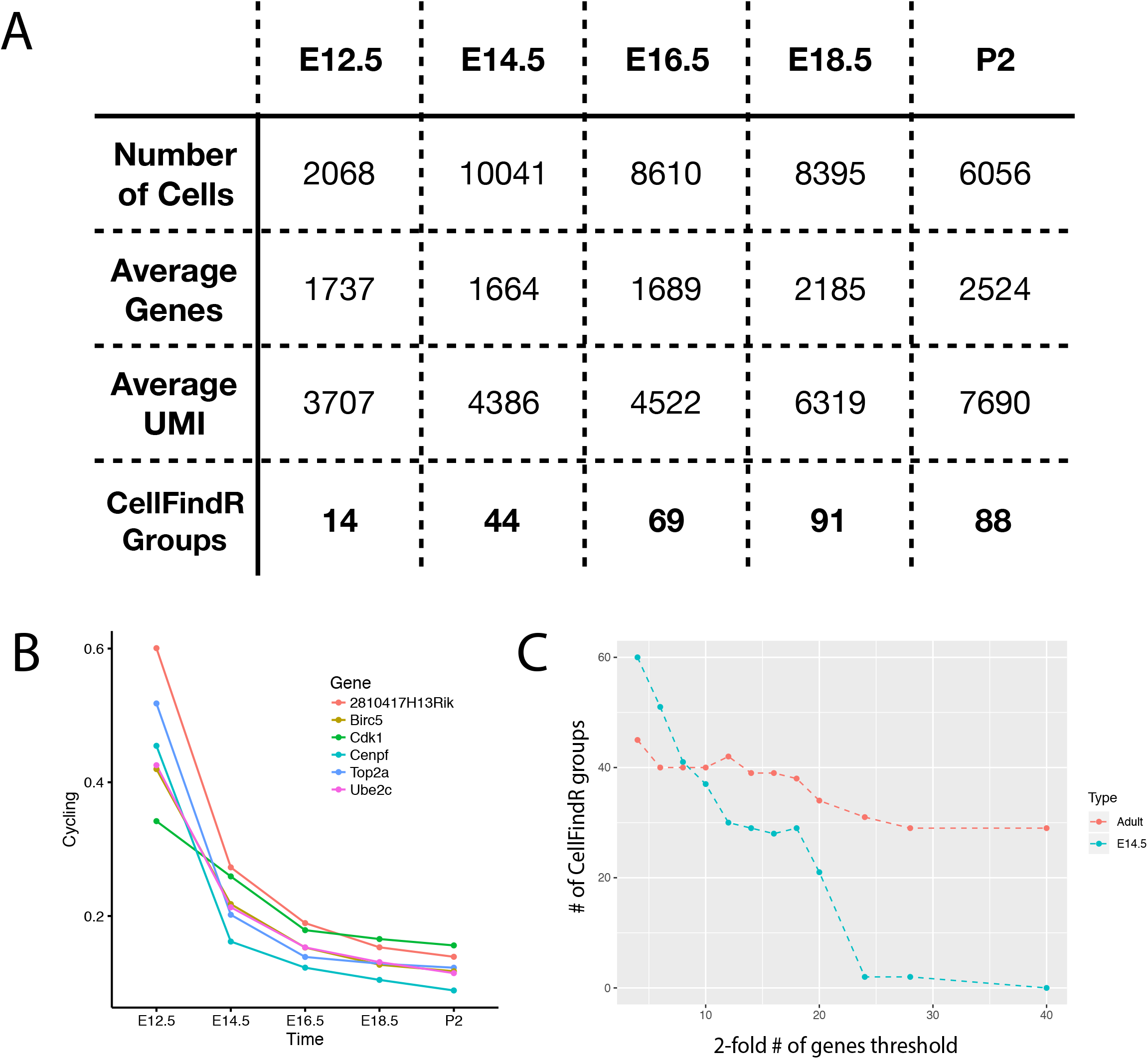
E12.5 to P2 mouse embryonic statistics and testing parameter threshold for CellFindR threshold. (A) Table of general statistics of each dataset from the mouse time course, showing the number of cells, distribution of the genes and number of CellFindR groups found. (B) Plotting the percentage of cells expressing known cycling genes through each dataset shows downtrending proliferation across time. (C) The threshold for number of genes that are two-fold differentially expressed are varied (x-axis) with the number of CellFindR output groups (y-axis) for both E14.5 and Adult *Epcam* sorted datatsets indicating stability of clustering.

**Figure S5:**
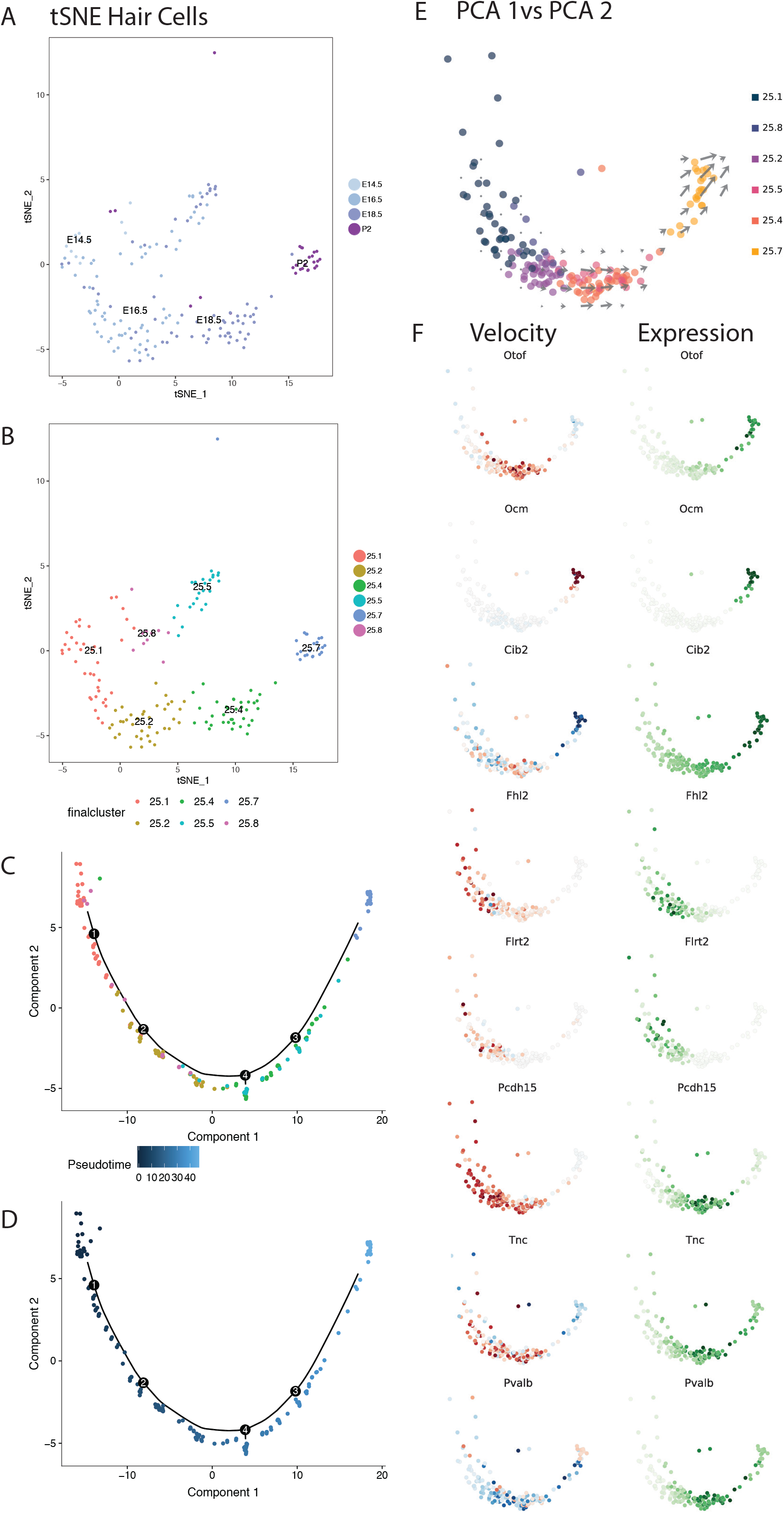
Trajectory analysis of hair cells with monocle and velocyto. tSNE representation of all mouse hair cells labeled with (A) the age of the cell and (B) with the CellFindR group definition. The trajectory generated by Monocle is plotted with (C) CellFindR groups and with the (D) calculated pseudotime, which confirm the validity of the Monocle model in estimating time based on our time course data. (E) PCA1 vs PCA2 representation of all mouse hair cells. (F) The RNA velocity plot: red represents induction, blue represents repression and expression plot (green) is shown for marker hair genes.

**Figure S6:**
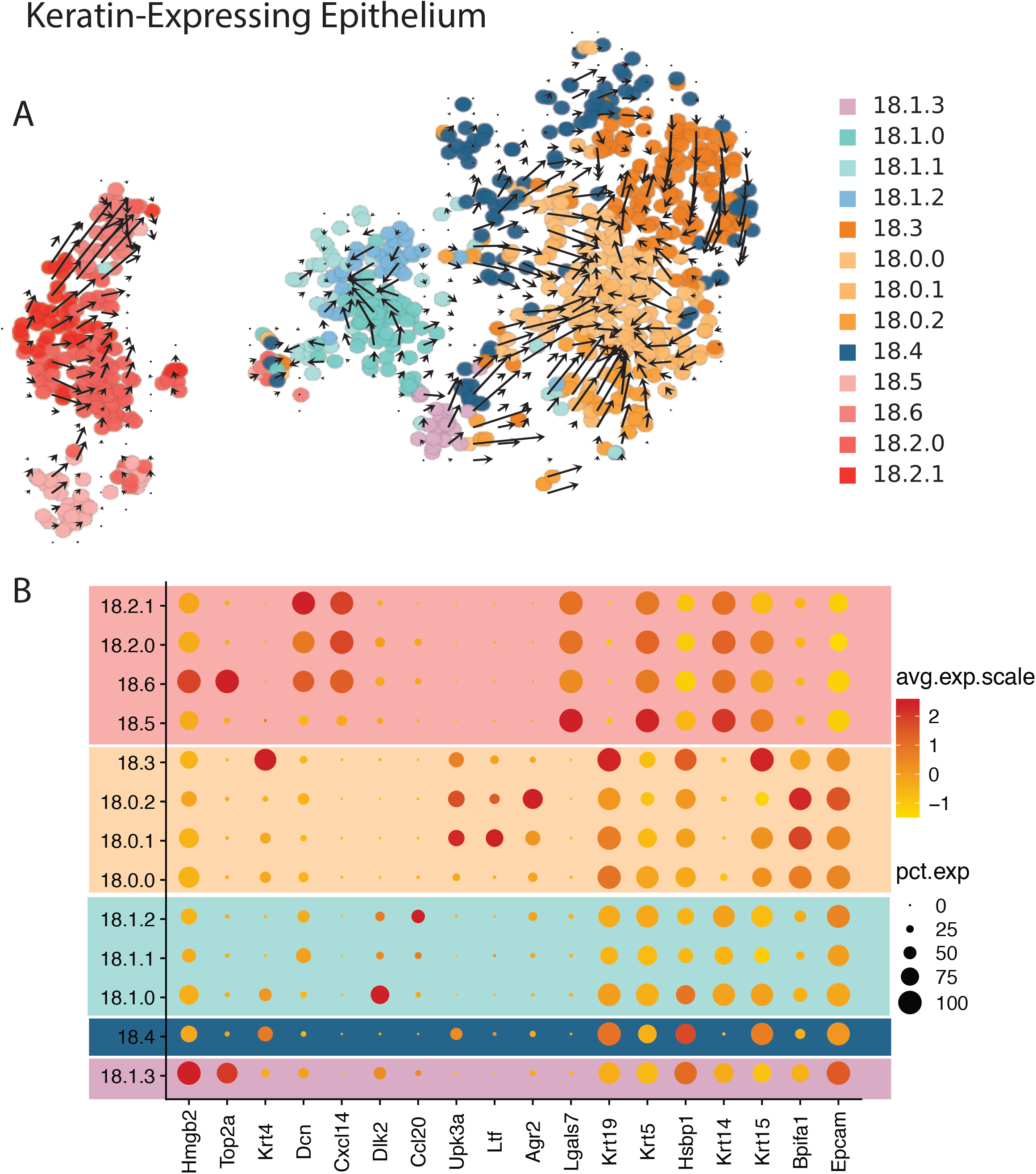
Keratin-expressing epithelium trajectory. (A) tSNE presentation with velocyto mapped trajectories. (B) Dotplot of differentially expressed genes for each group.

**Figure S7:**
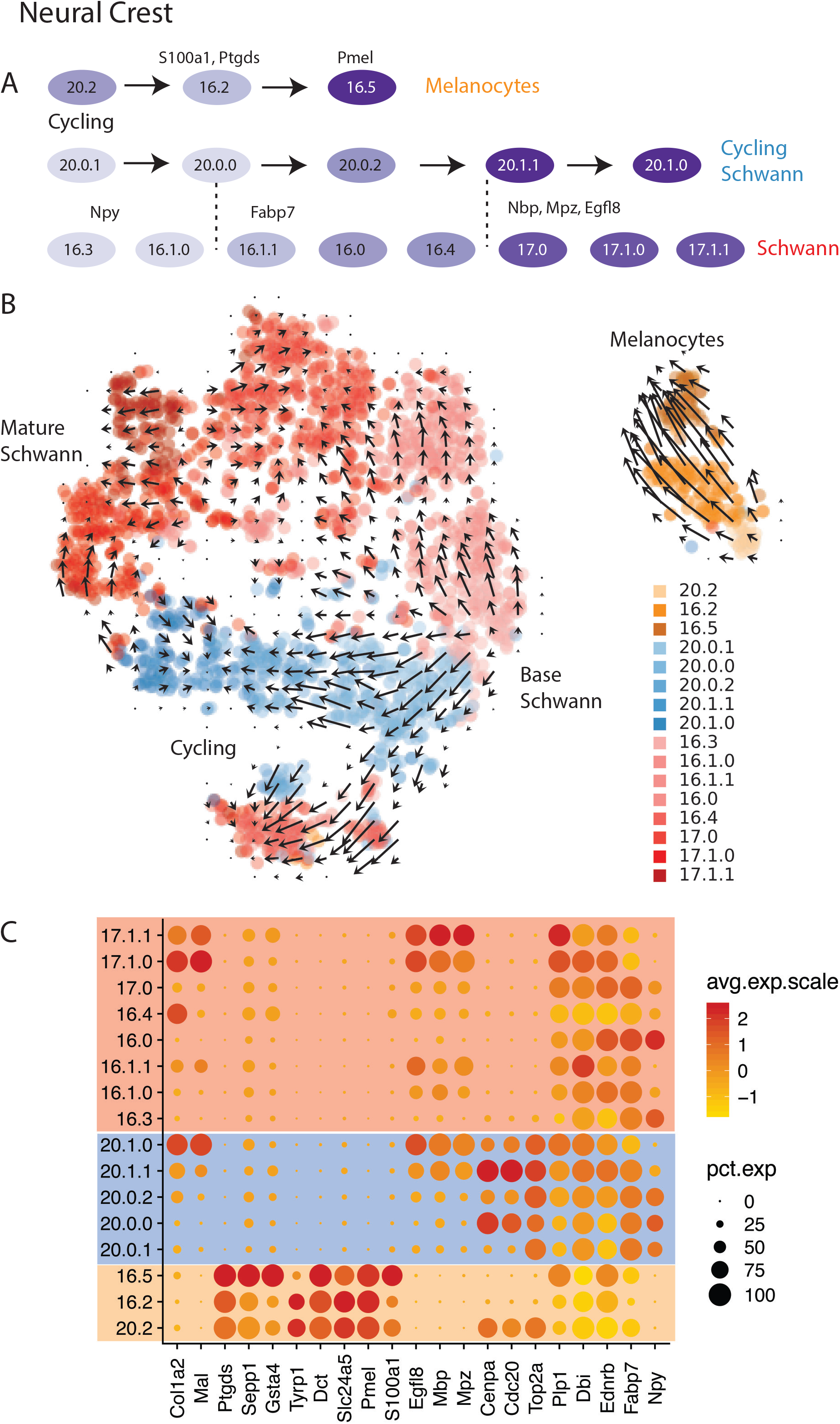
Neural crest and melanocyte trajectory. (A) Schematic lineage of CellFindR groups showing progression of cycling and non-cycling groups for the neural crest and a single linear trajectory of melanocytes. (B) UMAP representation with overlaid velocyto trajectories showing progression of schwann cell maturation and separate melanocyte group. (C) Dotplot of selected differentially expressed genes across the CellFindR groups showing unique time-correlated markers.

**Figure S8:**
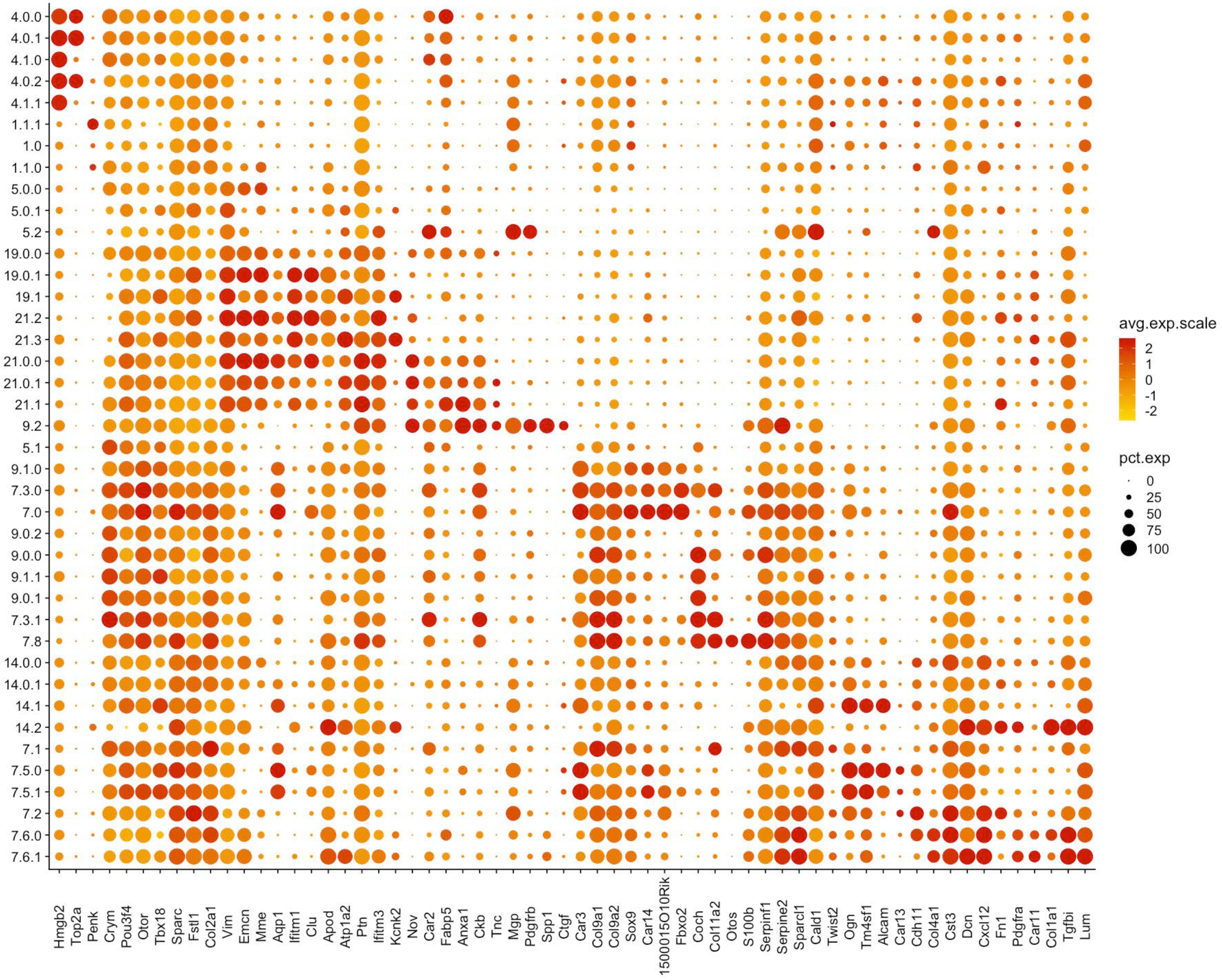
Mesenchymal CellFindR population dotplot. Expression of selected markers from all mesenchymal CellFindR fibrocyte groups are plotted.

**Figure S9:**
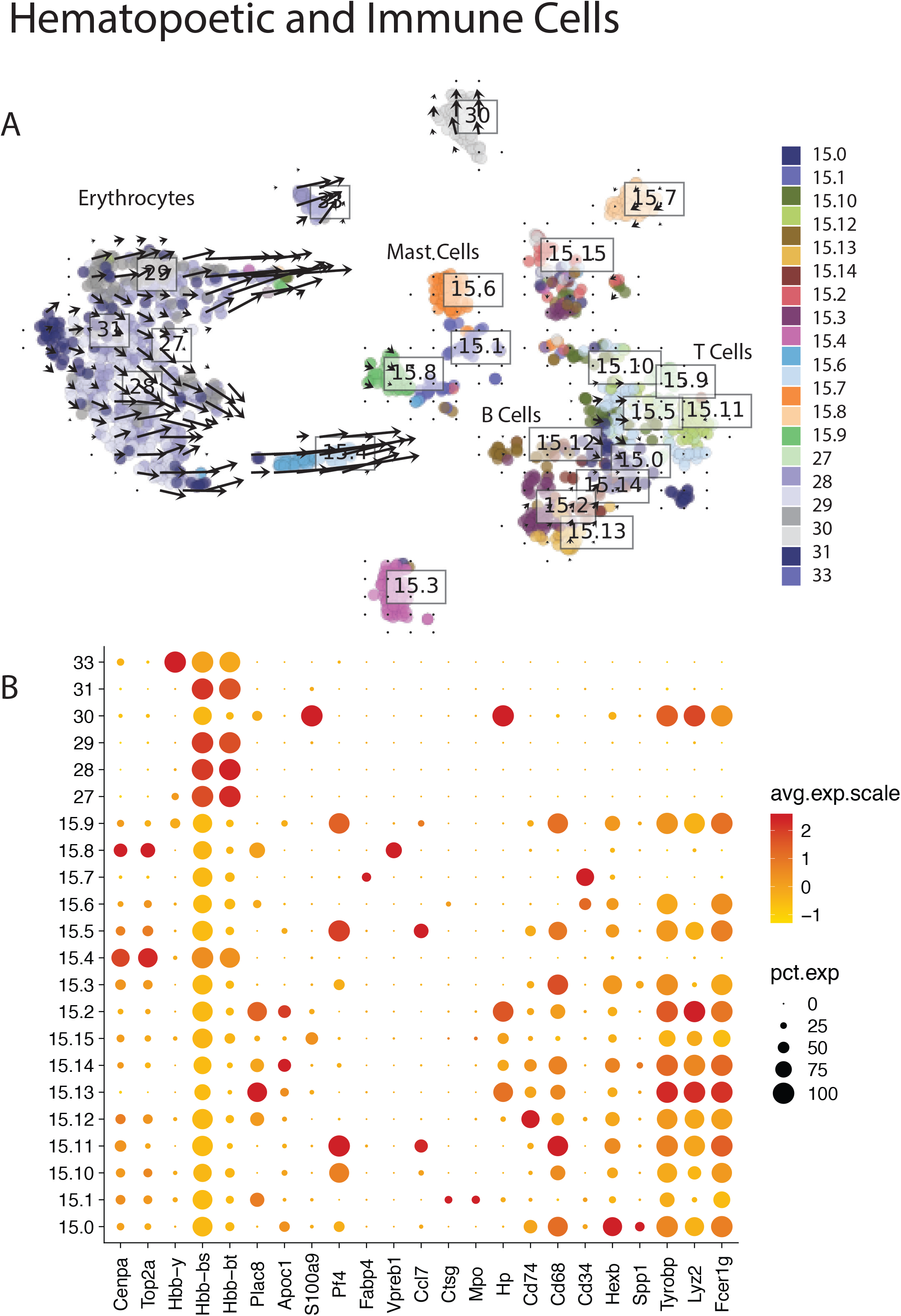
Hematopoietic and Immune cell populations. (A) tSNE representation with overlaid velocyto trajectory analysis shows most active changes in the erythrocyte maturation and few changes in the immune population. (B) dotplot showing marker genes for each of the separate CellFindR groups.

**Figure S10:**
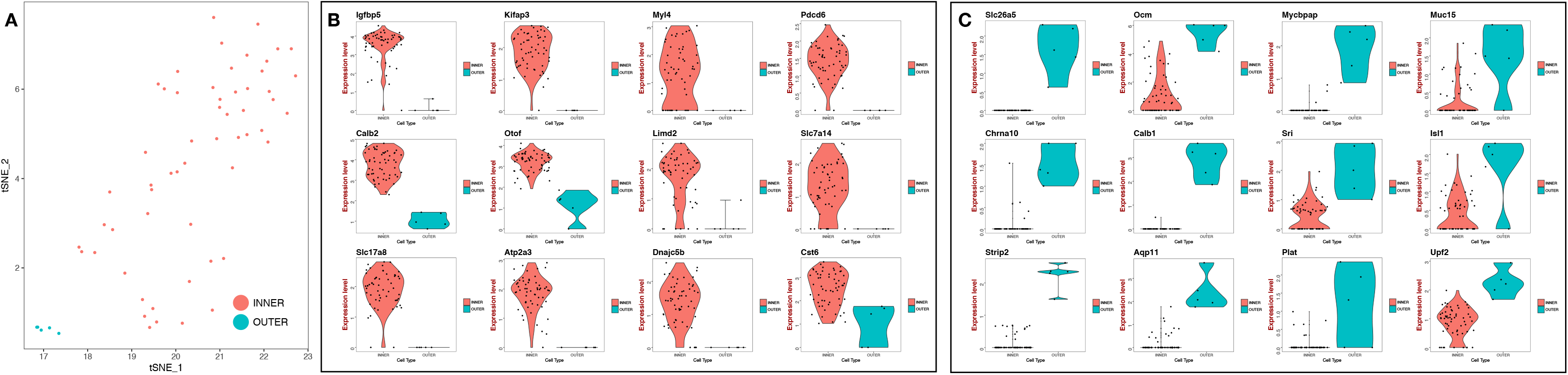
Inner and outer cochlear hair cells of the adult mouse. (A) tSNE representation of the adult mouse cochlea hair cells manually divided to inner and outer. Violin plots of log normalized expression of upregulated known and new markers for (B) inner and (C) outer hair cells.

**Figure S11:**
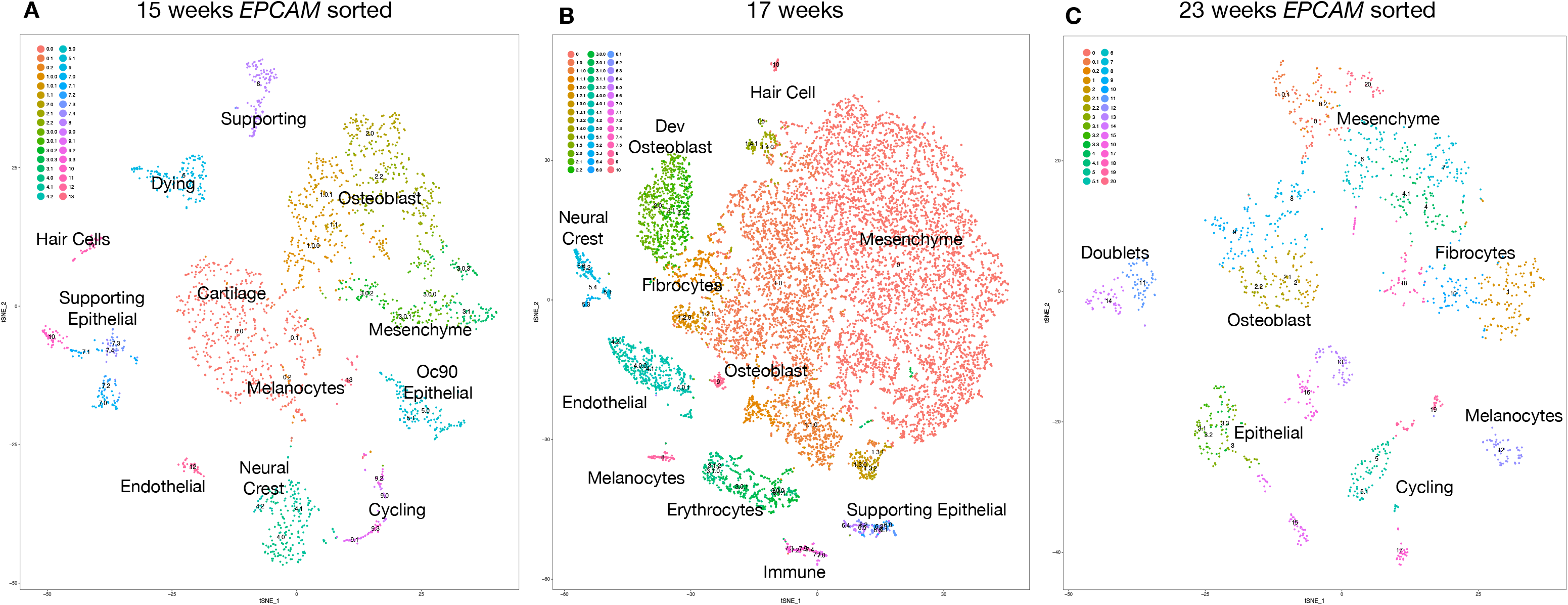
tSNE plots of fetal human (15, 17 and 23 weeks) cochlea. tSNE representation with labeled clusters for (A) 15 week human fetal *EPCAM* sorted, (B) 17 week human fetal, (C) 23 week human fetal datasets.

**Figure S12:**
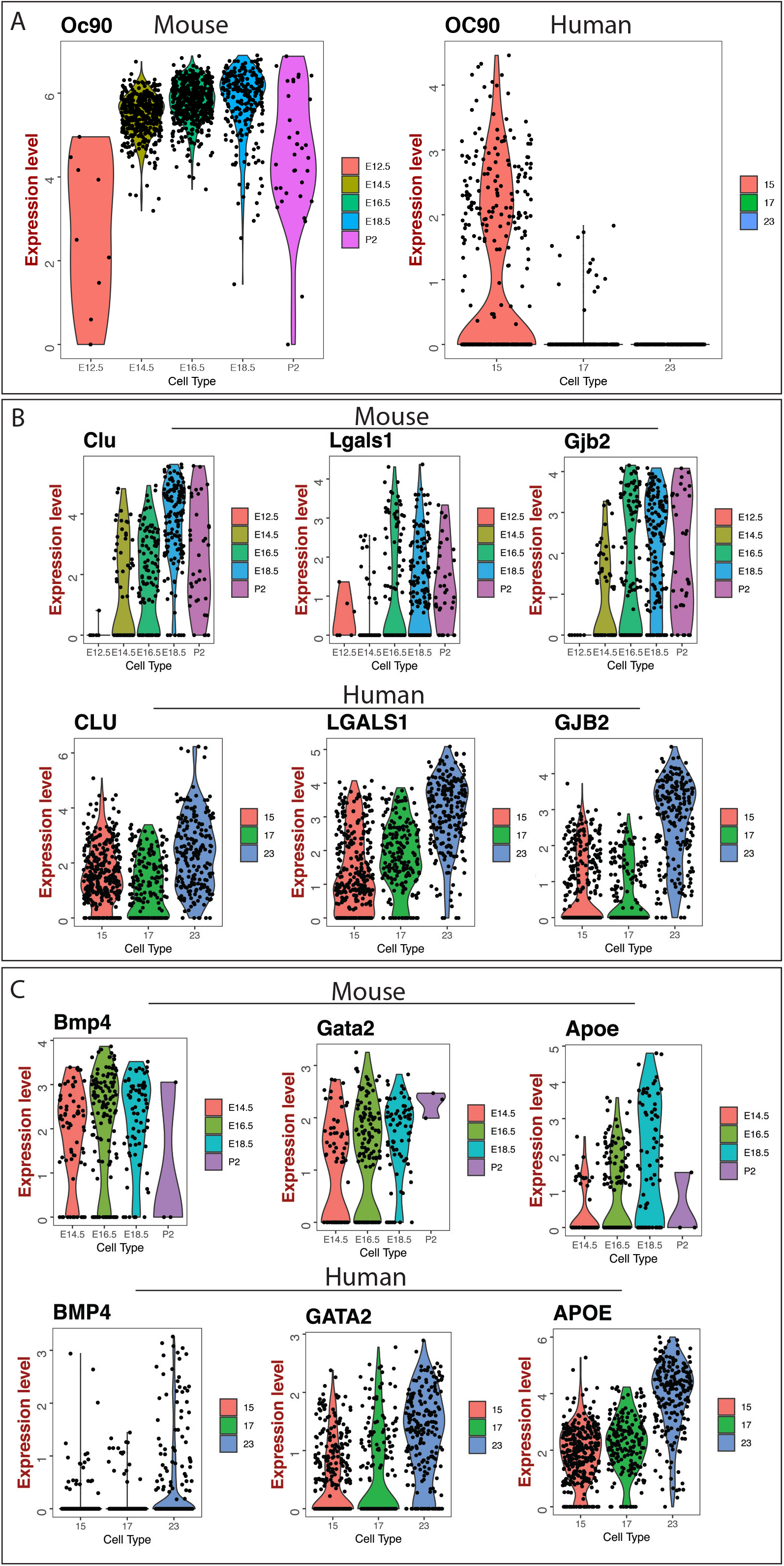
Comparing Expression of mouse and human markers across timepoints. (A) Oc90 expression in both mice and human epithelial subgroups. (B) Mouse and Human cochlear floor supporting epithelial markers across E12.5 to P2 mouse embryonic time vs 15 to 23 weeks in human fetal life. (C) outer sulcus markers between mouse and human.

**Figure S13:**
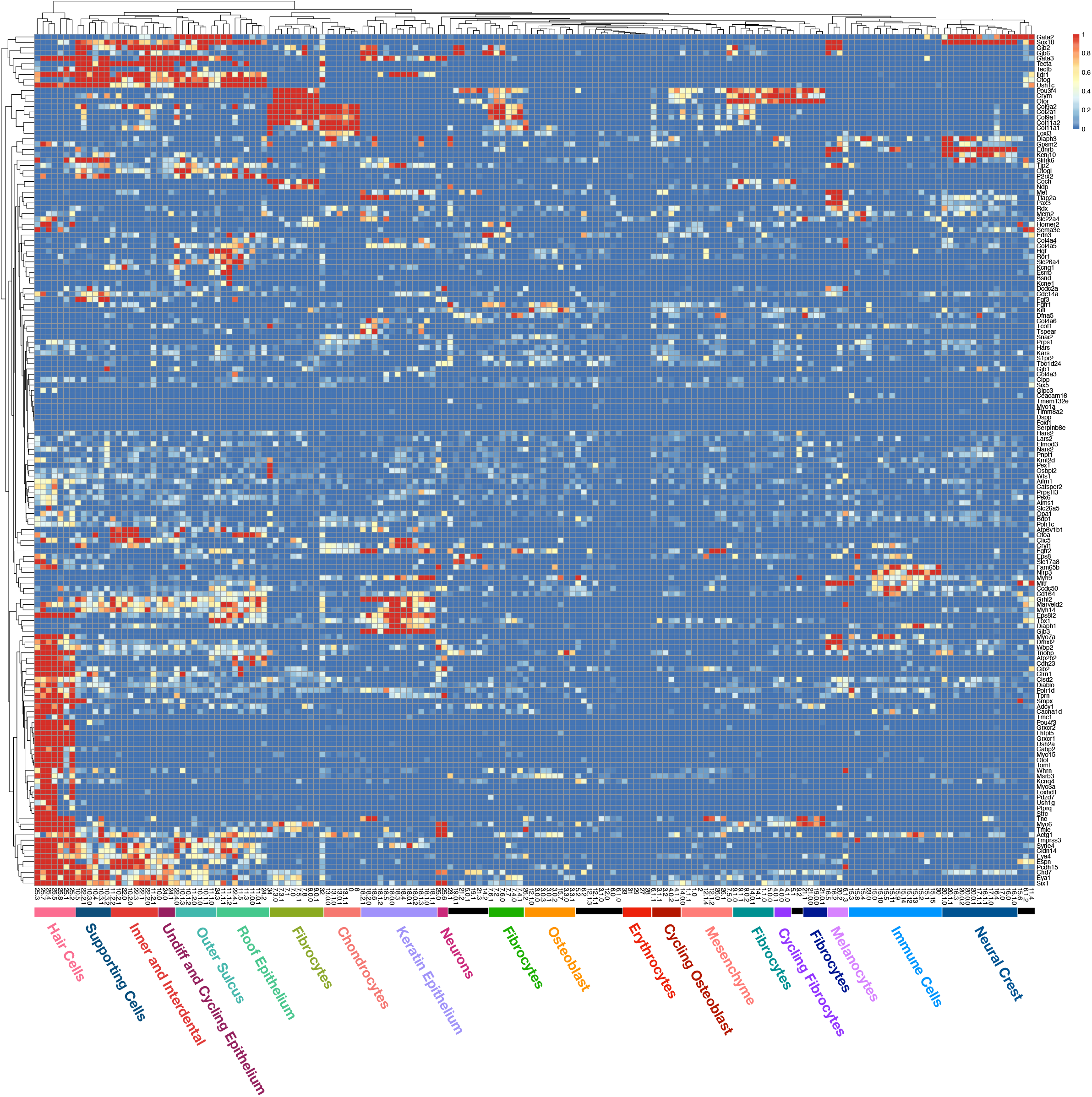
Deafness gene heatmap of the time course (E12.5-P2) embryonic mouse. Heatmap representing deafness genes (rows) and CellFindR groups of batched embryonic mouse dataset with clusters labeled by identity. Average expression levels are taken of each CellFindR group and binned to a maximum of 1. Both columns and rows of the heatmap are hierarchically clustered.

**Figure S14:**
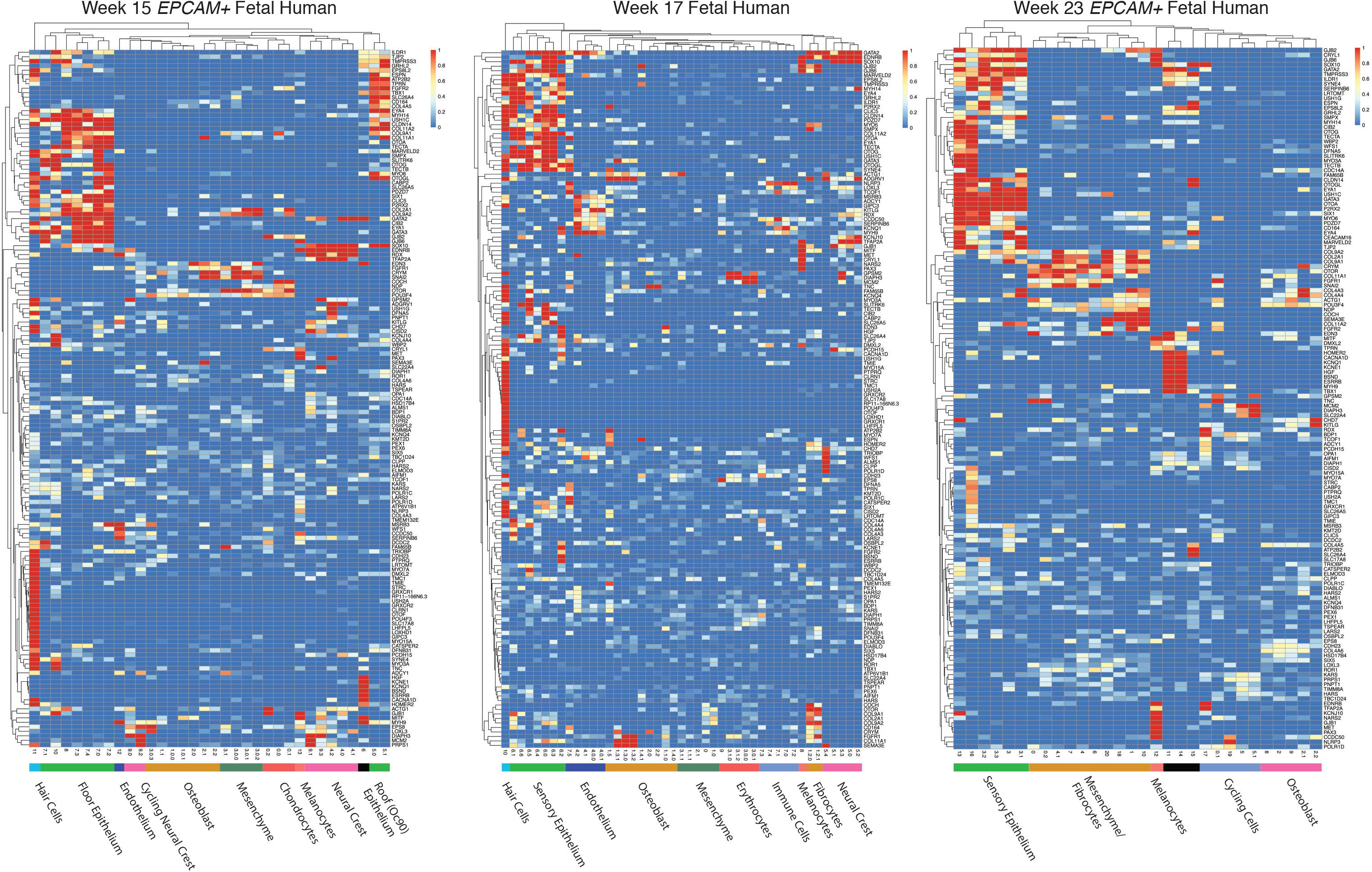
Deafness gene heatmap of the time course (15-23 week) fetal human. Heatmap representing deafness genes (rows) and CellFindR groups of with clusters labeled by identity. Average expression levels are taken of each CellFindR group and binned to a maximum of 1. Both columns and rows of the heatmap are hierarchically clustered. (A) 15 weeks EPCAM sorted, (B) 17 weeks, (C) 23 weeks EPCAM sorted.

## Methods

### Specimen collection for scRNA-seq analysis

All animal experiments were conducted in accordance with and approval of the UCSF Institutional Animal Care Use Committee. Single-cell data from developing cochleas were generated in two experiments. In the first pilot experiment, cochleas were harvested from E14.5 embryos from Swiss Webster mice. In the second, timed pregnant Swiss Webster females were purchased from Charles River Laboratories and cochleas were isolated from E12.5, E14.5, E16.5, and E18.5 embryos, as well as P2 pups. The tissue was minced with a scalpel, collected in PBS, and pelleted by centrifugation. It was dissociated in TrypLE Select (Life Technologies) at 37°C with trituration every five minutes. Once the tissue was dissociated (10-30 minutes), HBSS supplemented with 10% FBS and 2 mM EDTA was added and the suspension was passed through a 30 µm cell strainer. Cells were pelleted and then incubated with SYTOX Blue dead cell stain (Thermo Fisher Scientific) at a 1:1000 dilution for twenty minutes on ice. Cells were washed and then subjected to FACS to collect only live cells. Sorted cells were counted on a Moxi Flow (ORFLO Technologies) and resuspended at 1000 cells/µL in 0.04% BSA in PBS for single cell capture.

To identify the cochlear cell types in adult mice, two experiments were again conducted. In each experiment, cochlear soft tissue was dissected from 18 six-week-old Swiss Webster mice (nine male, nine female). After mincing with a scalpel and washing in PBS, the tissue was incubated in thermolysin (Sigma-Aldrich) at 37°C for 20 minutes. It was then transferred to Accumax (Innovative Cell Technologies) for 35 minutes, with trituration every 5-10 minutes. The dissociated tissue was diluted in PBS with 10% FBS and filtered through a 40 µm cell strainer. Cells were pelleted and dead cells and debris removed using the Dead Cell Removal Kit (Miltenyi Biotec) following the manufacturer’s instructions. When indicated, Epcam positive cells were next enriched in one fraction using CD326 (EpCAM) MicroBeads, mouse (Miltenyi Biotec). The resulting cells were counted and resuspended as indicated above.

Cochleas were harvested from gestational age: 15-week, 17-week, and 23-week post-mortem human fetuses with permission from the UCSF Human Research Protection Program Institutional Review Board. The soft tissue was dissected out, minced with a scalpel, and washed with PBS. The dissociation protocol described for the adult murine cochleas was followed: 20 minutes in thermolysin followed by 10 to 40 minutes in Accumax with trituration. Resulting cells were passed through a 40 µm cell strainer and pelleted, then red blood cells were lysed (Alfa Aesar), and the cells were processed using the Dead Cell Removal Kit. When indicated, MicroBeads containing anti-human CD326 (EpCAM) (Miltenyi Biotec) were used to enrich or deplete for EpCAM. Resulting cells were resuspended for single-cell capture as above.

### scRNAseq data processing

Single Cell 3’ Reagent Version 2 (v2) Kit and Chromium Controller (10x Genomics, CA, USA) were used as previously described (Zheng et al., 2017) to process cells before sequencing. Cellranger outputs from 2.0.0 were then run on the read files to output our count matrices. One verification dataset from E14.5 utilized the v1 kit. Data were analyzed using R and package Seurat (Butler et al., 2018). Cells were processed via the Seurat workflow to remove doublets and unwanted sources of variation. The basic filtering removed genes expressed in less than 3 cells and removed cells with less than 200 genes expressed with an upper limit cut of 7000 genes per cell. Principal component analysis was performed to reduce dimensionality on the scaled and log-normalized data matrix. We chose not to sort out mitochondrial genes. The first 10 PCA components were used as a basis for clustering for increased computational speed, and the Seurat R object was utilized for convenient manipulation and data transformation. Primary work was done in Seurat 1.4. For batching of the embryonic dataset, the merge function with regression over the datasets was conducted with Seurat’s algorithm. These sequences were run through the same sequencer in the same lane, thus, batching effects were potentially minimized.

### CellFindR pipeline

CellFindR relies on iterative Louvain clustering as implemented in the Seurat package. For an input Seurat object, CellFindR iterates through increasing resolution parameters for Louvain clustering (decreasing neighborhood sampling size) which increases the number of unique communities or clusters. It continuously tests for the condition that the generated clustering groups satisfy the condition that n-1 of the groups have ten genes that are two-fold differentially expressed when compared to the rest of the groups. These parameters were chosen based on the stability of the groupings and through examining the quality of groups. Both E14.5 and adult *Epcam* enriched dataset seem to have plateaus at ten genes (Fig S4c). Once these groups break this condition, each subsequent subgroup is then subjected to the same clustering conditions effective creating a branching tree of groups. CellFindR outputs a table for differential genes and their characteristics of each subgroup, a list of cluster identities for each cell labeled by barcode, and a list of the resolutions chosen for each clustering step. CellFindR requires R running at 3.4.4 with version 1.4 of Seurat as well as other packages: matrix, ggplot, pheatmaps, RColorBrewer. Github source code can be found at: https://github.com/kevyu27/CellFindR.

### Tissue processing for RNAscope

For staining: 40 cochleas were collected from 20 wildtype FVB/NJ mice (Charles River). Before dissection, mice were transcardially perfused, first with RNase free PBS, then with 4% paraformaldehyde in RNase free PBS. All solutions used were RNase free to prevent RNA degradation in preparation for RNAscope® protocol. After dissection, cochleas were fixed for 24 hours at 4°C in 4% paraformaldehyde. Cochleae were then decalcified in 5% EDTA for three to four days at 4°C, with daily solution changes. In order to cryoprotect samples, cochleae were rocked in 30% sucrose for one hour at room temperature. Cochleae were embedded in O.C.T. freezing medium (Fisher Scientific), kept at 4°C for at least one hour to allow for complete equilibration of tissue in embedding medium, and then frozen overnight at −80°C. 12µm thick serial sections parallel to the modiolus were taken using a Cryostar™ NX70 Cryostat (ThermoFisher Scientific). Sections were placed on Fisherbrand™ Superfrost™ Plus Slides (Fisher Scientific) and frozen at −80°C.

For paraffin sections, fixed and decalcified cochleae were dehydrated in graded concentrations of ethanol (30%, 50%, 70% 90%, 100%), then cleared in Histo-clear II (Electron Microscopy Sciences) at room temperature. Cochleae were paraffin infused overnight at 62°. Then cochleae were embedded in paraffin blocks and cut into 7 µm thick sections and mounted onto Fisherbrand™ Superfrost™ Plus Slides (Fisher Scientific) and baked at 37°C overnight. For both paraffin and optimal cutting temperature compound (OCT) sections, two representative slides were stained with Hematoxylin 1 and Eosin-Y (Fisher Scientific) and imaged with a standard bright-field microscope to confirm tissue structure and quality of sections.

### RNAscope validation of chosen markers in OCT sections

Using data from CellFindR, gene markers were chosen for various clusters of interest based on highest average differentially expressed genes for each cluster. Markers were chosen based on cluster specificity and high level of expression. RNAscope® probes for these markers were purchased from Advanced Cell Diagnostics. RNA *in situ* hybridization of the cochlea was performed using the RNAscope® Multiplex Fluorescent Reagent Kit v2 (Advanced Cell Diagnostics). In order to maintain tissue integrity throughout the staining protocol, slides were post-fixed in 4% paraformaldehyde in RNase free PBS, washed twice in PBS, and baked at 60°C for 30 minutes. To permeabilize tissues for target probe hybridization, sections were incubated in hydrogen peroxide for 10 minutes at room temperature, washed in PBS, then incubated in RNAscope® Protease Plus for 30 minutes. Slides were then washed in distilled water and incubated with target probes for two hours. After target probe hybridization, multiple signal amplification molecules were hybridized to target probes as per the RNAscope® Multiplex Fluorescent v2 Assay protocol. Slides were imaged on a Leica DM6 B microscope. Sections probed for gene markers were compared to both negative and positive control slides to validate probe signals.

### RNAscope validation of chosen markers in paraffin sections

Paraffin slides were baked at 60°C for one hour, then deparaffinized in two changes of Histo-clear II (Electron Microscopy Sciences) for five minutes, then immersed in two changes of 100% ethanol for two minutes, then baked at 60°C for five minutes. Sections were incubated in hydrogen peroxide for 10 minutes to permeabilize the tissue. Target retrieval was performed by boiling slides at 98°C in RNAscope® 1X Target Retrieval Reagents for 15 minutes. Then slides were incubated in RNAscope® Protease Plus for 30 minutes at 40°C. Slides were then washed in distilled water and incubated with target probes for two hours. After target probe hybridization, multiple signal amplification molecules were hybridized to target probes as per the RNAscope® Multiplex Fluorescent v2 Assay protocol. Slides were imaged on a Leica DM6 B microscope. Sections probed for gene markers were compared to both negative and positive control slides to validate probe signals.

### RNAscope validation of chosen markers in wholemount Organ of Corti

Whole-mount RNAscope® was performed according to the protocol established by the Fritzsch lab at University of Iowa (Kersigo et al., 2018). Micro-dissected pieces were dehydrated in a graded MeOH/PBT series (50%, 75%), then incubated in 100% MeOH for either five minutes or stored overnight at −20°C. Tissue was rehydrated in the same graded MeOH series in reverse, then washed for five minutes three times in PBT. Then tissue was incubated in RNAscope® Protease III for 20 minutes. Tissue was washed in PBT then incubated with selected probes overnight at 40°C. After target probe hybridization, tissue was washed in 0.2x SSC and incubated in multiple signal amplification molecules as per the RNAscope® Multiplex Fluorescent v2 Assay protocol. After amplification molecule and fluorophore hybridization was completed, tissue was incubated in Dapi for 15 minutes and mounted onto glass slides using Prolong Gold Anti-fade Mountant and imaged using a Nikon C2 MacroConfocal.

### Deafness gene curation

Genes from the Hereditary Hearing Loss Homepage, University of Iowa, OtoSCOPE, for genetic testing were pulled together (Azaiez et al., 2018). Manipulations of data was conducted in R. The average expression of each deafness gene in a CellFindR subgroup of cells were calculated. The upper limit ceiling was set at a log-expression normalized threshold of 1 and this was plotted onto the heatmap through pheatmap R package with color shading from the RColorBrewer package.

For the combined mouse heatmap, we generalized the expression of a particular deafness gene at each time point to the CellFindR subgroup with the highest expression. For example, at a given timepoint, for gene A, we find the maximum expression level to be in subgroup 1, and thus will use that as the gene expression. This value was collated for each gene and each time point. Genes were ordered by hierarchical clustering and then sorted chronologically based on the weight contribution from each time point to create a sorted heatmap of expression from E12.5 to Adult.

### Correlation matrices and heatmaps

The average expression matrix was generated using function avg_plot documented in CellFindR. This function takes the scaled matrix and averages all the expression levels of each gene in all the cellular subgroups. Once this is generated, the intersection function in CellFindR was used to identify the top 100 differentially expressed genes in the two datasets and then the number of matching genes, hence the intersection of the differentially expressed genes were generated into a numeric heatmap. For those comparing mouse to human, a table was created organizing homology of human and mouse genes to a common label number. The top 100 genes were translated to the common label number and then the number of matches were counted to create a similar numeric heatmap.

### Statistical Analysis

For differential gene analysis of groups, the ‘bimod’, likelihood-ratio test for single cell gene expression was used to determine significant genes (McDavid et al., 2013). The p-values are then renormalized by Bonferroni adjustment to correct for the multiple comparisons. These methods were adapted from Seurat.

### Monocle Analysis

In order to infer cellular trajectories, we used the Monocle package (2.8) in R (Trapnell et al., 2014). Seurat objects were converted to CellDataSet objects compatible with Monocle and were analyzed using their downstream analyses with normalization and filtering through their estimating size factor and dispersion functions. Filtering steps were conserved from pseudotime, trajectory analysis, and differential gene calculations were conducted along the branches following methods described in their vignette.

### Velocyto Analysis

We performed RNA velocity-based trajectory inference analysis using the Python implementation of Velocyto (v0.17) (La Manno et al., 2018). First, BAM files were generated using Cell Ranger, with reads aligned against standard mouse (mm10) or human reference genomes (hg19 or GRCh38). The velocyto.py command line interface was used to process the BAM files into loom files with gene expression counts annotated as unspliced, spliced, or ambiguous. The loompy package (v2.0.17) was used to merge loom files where appropriate, and to annotate metadata as well as assign CellFindR group identities based on individual cell barcodes. Scanpy (v1.4.1) was used for standard pre-processing as performed in Seurat, including log-normalization and zeroing of expression levels (Wolf et al., 2018). Scanpy was also used for data visualization including generation of UMAP and tSNE plots. RNA velocity estimates were calculated on related cell populations of interest using filtering and analysis parameters determined in the manner described in La Manno et al 2018.

---

**Table 1: Mouse E14.5 Cochlear Epithelial Floor Expression and Differential Expression.** Output from CellFindR for E14.5 cochlear epithelium floor with columns representing log-normalized: mean, standard deviation, average differential expression and corrected p-values for each CellFindR group. Rows represent each measured gene. Blank represents non-significant value.

**Table 2: Mouse Embryonic Epithelium Floor Expression and Differential Expression.** Output from CellFindR for batched E12.5 to P2 epithelium floor with columns representing log-normalized: mean, standard deviation, average differential expression and corrected p-values for each CellFindR group. Rows represent each measured gene. Blank represents non-significant value..

**Table 3: Summary of scRNAseq experiments**. All experiments labeled with their qualities metrics and cell counts.

